# The Core Subunit NDUFS2 in Mitochondrial Complex I is Critical to Oxygen Sensing in Human Ductus Arteriosus Smooth Muscle Cells

**DOI:** 10.1101/2025.07.08.663799

**Authors:** Rachel E.T. Bentley, Kimberly J. Dunham-Snary, Ashley Y. Martin, Jeffrey D. Mewburn, Asish Dasgupta, Kuang-Hueih Chen, Benjamin P. Ott, Tanvi Nandani, Bernard Thébaud, Mark K. Friedberg, Charles C.T. Hindmarch, Stephen L. Archer

**Affiliations:** Department of Medicine, Queen’s University, Kingston, Ontario, Canada; Department of Biomedical and Molecular Sciences, Queen’s University, Kingston, Ontario, Canada; Queen’s Cardiopulmonary Unit, Translational Institute of Medicine (TIME), Department of Medicine, Queen’s University, Kingston, Ontario, Canada; Regenerative Medicine Program, Ottawa Hospital Research Institute, Ottawa, Ontario, Canada; Department of Cellular and Molecular Medicine, University of Ottawa, Ottawa, Ontario, Canada; Neonatology, Department of Pediatrics, Children’s Hospital of Eastern Ontario (CHEO) and CHEO Research Institute, Ottawa, Ontario, Canada; Division of Cardiology, The Labatt Family Heart Center, Hospital for Sick Children and University of Toronto, Toronto, Ontario, Canada

## Abstract

**Background:** Mitochondria in ductus arteriosus smooth muscle cells (DASMCs) are oxygen sensors triggering vasoconstriction at birth; however, the oxygen sensing mechanisms are incompletely understood. Given the conserved role of mitochondrial Complex I subunit NDUFS2 in other oxygen-sensing tissues, we examined its role in DASMC oxygen sensing, comparing it to other Complex I subunits (NDUFS1 and NDUFS7) and putative O_2_-sensor subunits (UQCRFS1 and COX4I2).

**Methods:** Human DASMCs were grown in hypoxia (pO_2_=41mmHg). Oxygen responsiveness was assessed, measuring changes in intracellular calcium, [Ca^2+^]_i_, cell length, and mitochondrial reactive oxygen species (mROS). DASMCs were treated for 48-hours with control siRNA versus siRNA targeting NDUFS2, NDUFS1, NDUFS7, UQCRFS1, or COX4I2. qPCR and immunoblotting confirmed knockdown. 3’RNA sequencing assessed transcriptional changes following siRNA.

**Results:** Oxygen increased mitochondrial fission, [Ca^2+^]_i_, and constricted DASMCs. 48-hours post-treatment, siNDUFS2 selectively depressed oxygen-induced increase in [Ca^2+^]_i_ (siControl +18.6±2.3%, siNDUFS2 +5.5±1.5%, p<0.0001), DASMC shortening (from 18.4±1.1% to 8.9±0.8%, p<0.0001), and mROS (+24±4.9% untreated, -6.6±5.4% post-siNDUFS2, p<0.0001), without altering the KCl response or depressing respiration. The mitochondrial antioxidant MitoTEMPO reduced mROS (2.9±4.5%, p=0.001) and attenuated oxygen-induced DASMC shortening (8.4±0.9%, p=0.0003). Transcriptomics revealed unique changes in mitochondrial pathways post siNDUFS2.

**Conclusions:** NDUFS2 regulates mROS and is a mitochondrial oxygen sensor in human DASMCs.

**Impact:**

- We demonstrated a unique role of Complex I subunit NDUFS2 (NADH:Ubiquinone Oxidoreductase Core Subunit S2), amongst putative oxygen-sensing electron transport chain subunits, in the responsiveness of human ductus arteriosus (DA) smooth muscle cells (DASMCs) to oxygen.
- NDUFS2 knockdown inhibited oxygen-induced DASMC constriction and generation of mitochondrial reactive oxygen species, at a timepoint prior to inhibition of mitochondrial respiration and without inhibition of KCl-induced constriction.
- While mitochondria are known DA oxygen sensors, this work identifies NDUFS2 as a molecular mediator of human DA oxygen sensing within the mitochondria, enhancing our understanding of a vital physiologic phenomenon and providing a novel potential therapeutic target to modulate ductal patency.

## 1. Introduction

The transition of the circulatory system from placental oxygenation in the fetus to an air-breathing neonate relies on the response of the ductus arteriosus (DA), a fetal vessel connecting the pulmonary artery and aorta. The acute functional constriction of the DA occurs in the hours after birth, triggered by the increase in arterial partial pressure of oxygen (pO_2_); this not only stops blood flow through the DA, terminating right-to-left shunting^1^, but is a necessary precondition for permanent anatomical DA closure^2^. One of the most common complications of preterm birth, persistently patent ductus arteriosus (PDA), occurs when the DA fails to close^3,4^. PDA occurs in 55-70% of neonates with birth weight under 1,000g and often causes hemodynamic compromise^5^, with a prolonged postnatal left-to-right shunt associated with heart failure and other complications^6^. Although our understanding of the DA’s oxygen response has increased, particularly our ability to enhance DA closure, using cyclooxygenase inhibitors like ibuprofen or meclofenamate^7^, or using prostaglandins to maintain patency in ductus-dependent congenital heart diseases (DD-CHD)^8^, the molecular identity of the DA oxygen sensor within the mitochondria remains elusive and clinically relevant^9,10^.

Despite the important role of the endothelium in DA closure, multiple groups have demonstrated endothelium-independent DA oxygen responses, suggesting that oxygen sensing primarily occurs in the DA smooth muscle cells (DASMCs)^11,12^. Increased oxygenation and the onset of ventilation initiate a signaling cascade within DASMCs, resulting in constriction. This cascade is initiated by increased mitochondrial-derived reactive oxygen species (mROS) generation, released during the process of mitochondrial fission^13^. Mitochondrial ROS trigger a redox-mediated inhibition of voltage-gated potassium channels (Kv)^13^. Kv inhibition depolarizes the DASMCs, increasing the intracellular calcium concentration and inducing vasoconstriction (**Figure 1A-B**)^14^. This rapid ionic mechanism of DA constriction, occurring within seconds to minutes of birth, is ultimately reinforced by release of endothelial derived vasoconstrictors, like endothelin 1, and withdrawal of vasodilator prostaglandins, both of which are regulated at term to promote DA constriction^15,16^.

**Figure 1:**
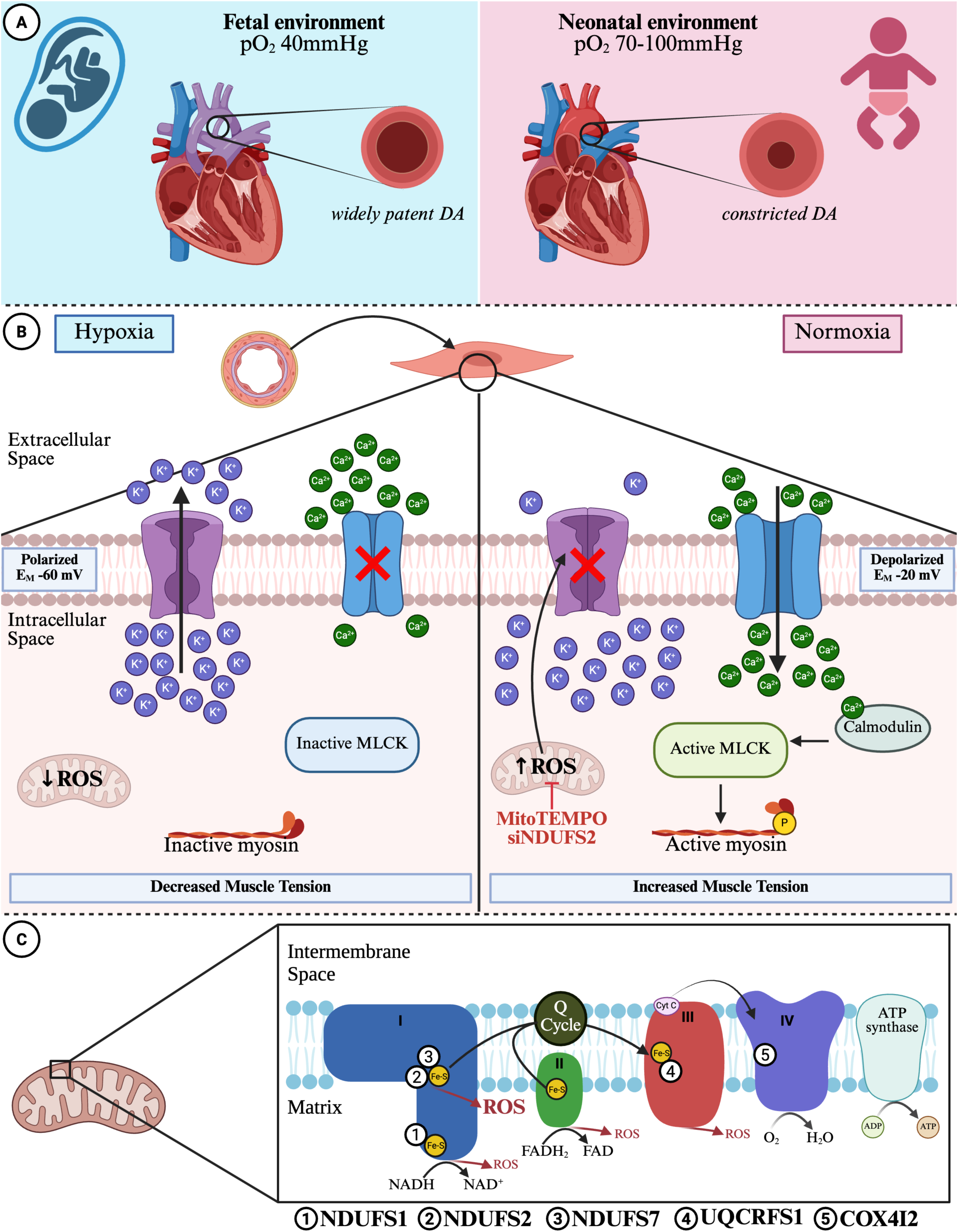
Ductus Arteriosus Smooth Muscle Cell Oxygen Sensing and Experimental Design. **A**) The fetal (left) hypoxic arterial partial pressure of oxygen (pO_2_, ∼40 mmHg) maintains a patent ductus arteriosus (DA), diverting placentally oxygenated blood to the aorta. The neonatal (right) rise in pO_2_ (70-100 mmHg) causes DA constriction. **B**) In DA smooth muscle cells (DASMCs), in hypoxia (left), there is low mitochondrial generation of reactive oxygen species (ROS), maintaining a polarized resting membrane potential (E_M_) and low concentrations of intracellular calcium [Ca^2+^]_i_ (voltage-gated potassium channels = purple, L-type calcium channels = blue). Under normoxic conditions (right), increased mitochondrial ROS generation promotes influx of calcium and activation of myosin to increase DASMC tension. The mitochondrial-targeted antioxidant, MitoTEMPO, and knockdown of NDUFS2 (siNDUFS2) both inhibit the increase in ROS. **C**) To examine the importance of putative mitochondrial O_2_- sensor proteins we selectively knocked down **Complex I subunits: 1**) NADH:Ubiquinone oxidoreductase core subunit S1 (NDUFS1), located near the NADH binding site. **2**) NADH:Ubiquinone oxidoreductase core subunit S2 (NDUFS2), at the ubiquinone binding site. **3**) NADH:Ubiquinone oxidoreductase core subunit S7 (NDUFS7), also at the ubiquinone binding site. **Complex III subunit: 4**) Ubiquinol-cytochrome c reductase, Rieske iron-sulfur polypeptide 1 (UQCRFS1), at the Rieske iron-sulfur cluster. **Complex IV subunit: 5**) Cytochrome c oxidase subunit 4I2 (COX4I2). Created in BioRender. Bentley, R. (2025) https://BioRender.com/o096p30

The role of mitochondria as oxygen sensors in specialized oxygen sensitive tissues, including the DA, pulmonary arteries, and carotid bodies, is well reported^17^. This collection of highly oxygen sensitive tissues that respond rapidly to physiologic changes in oxygenation is termed the homeostatic oxygen sensing system (HOSS) and functions to optimize systemic oxygen delivery^17^. The constriction of pulmonary arteries (PA) to hypoxia, a process known as hypoxic pulmonary vasoconstriction (HPV), enables ventilation-perfusion matching to optimize systemic oxygenation^18^, while hypoxic stimulation of the carotid body type 1 cells leads to neurosecretion and activation of the brainstem respiratory center^19^, which increases ventilation and improves systemic oxygenation by increasing oxygen uptake from the environment^17^. There are conserved oxygen sensing mechanisms in all HOSS tissues, with oxygen-sensitive potassium channels mediating the responses of the DA, PA, and carotid body^14,19,20^. Mitochondrial heterogeneity plays a central role in the opposing oxygen responses of the DA and PA, the DA constricting in response to normoxia at birth versus the PA which undergoes vasodilatation. Inhibitors of ETC Complex I (rotenone) and Complex III (antimycin A) recapitulate the opposing response to hypoxia in DA and PA vascular rings and isolated SMC, relaxing the DA and constricting the adult PA^14,21,22^. This heterogeneity in mitochondrial signaling in response to changes in oxygen contrasts the downstream ionic effector pathway which are similar in DA versus PA, with Kv channel inhibitors causing concordant vasoconstriction, and L-type calcium channel inhibitors causing concordant vasodilation in both DA and PA^14,23^.

Complex I is a major site of mROS generation^24^, with recent evidence supporting a key role for Complex I-specific mROS generation in mediating oxygen-induced DA constriction^25^. Specific subunits of ETC Complex I mediate oxygen responses of several HOSS tissues. Fernandez-Aguera et al. and our group have demonstrated that the Complex I subunit at the ubiquinone binding site (I_Q_), NDUFS2 (NADH:ubiquinone oxidoreductase core subunit S2), is necessary for oxygen sensing of the carotid body glomus cells^26^ and adult pulmonary artery smooth muscle cells^27^, respectively; however NDUFS2’s role in DASMCs is uncertain. A unique role for NDUFS2 in constriction to oxygen in human DA is suggested by the RNAseq profile of human DASMCs grown in hypoxia and then exposed to 4 days of normoxia, mimicking the fetal and neonatal oxygen conditions, respectively^28,29^. *NDUFS2* was uniquely downregulated by oxygen exposure amongst differentially expressed genes in Complex I, with expression of the other subunit I genes being upregulated^28^.

Here we elucidate the contribution of various candidate ETC subunits to DA oxygen sensing by selectively knocking down NDUFS2 and other subunits using silencing RNA (siRNA) and examining the functional consequences (**Figure 1C**). We examined the contribution of other putative mitochondrial oxygen sensor subunits, which have largely been identified in adult PASMCs and carotid bodies^30^, but have not previously been assessed in fetal arteries, including the DA. These additional oxygen sensor candidate subunits studied include: the Rieske iron sulfur protein in complex III, encoded by *UQCRFS1* (Ubiquinol-Cytochrome C Reductase, Rieske Iron-Sulfur Polypeptide 1)^31^, and Complex IV subunit COX4I2 (Cytochrome C Oxidase Subunit 4I2)^32^. We also investigated NDUFS7, a Complex I subunit in the I_Q_ site adjacent to NDUFS2, and NDUFS1, a Complex I subunit at the NADH binding site. Comparing effects of NDUFS2 knockdown to knockdown of subunits elsewhere within Complex I allowed us to determine if flow of electrons from NADH through Complex I (NDUFS1) or general inhibition of the ubiquinone binding site (NDUFS7) explain the inhibitory effects seen with siNDUFS2. Our findings support a unique role for NDUFS2 as an O_2_ sensor in human DASMCs (hDASMCs).

## 2. Materials & Methods

Comprehensive descriptions of materials and methodology are included in the supplementary materials. Queen’s University ethics approval was obtained from the Health Sciences and Affiliated Teaching Hospitals Research Ethics Board (HSREB) for the continued use of patient-derived human DASMC lines (TRAQ#6007784). Primary human DASMCs were used for all experiments. The limited demographic information available for the donors is provided in in **Table S2,** and the specific cells used for each corresponding result are listed in **Table S3**.

## 3. Results

Ten primary hDASMC cell lines, each with documented responsiveness to oxygen^28^, were used for all *in vitro* studies. Acute oxygen response was confirmed in all cell lines, quantified as an increase in mitochondrial fission, increased mitochondrial generation of ROS, increased intracellular calcium, and a decrease in cell length following 10-20 minutes exposure to oxygen, discussed in more detail below. The demographics of the 10 infants from whom the DASMCs were isolated are summarized in **Table S2**.

### 3.1. Knockdown Optimization

siRNA targeting *NDUFS1*, *UQCRFS1*, and *COX4I2* were selected based on prior work in pulmonary artery SMC (PASMCs)^27^, with the *COX4I2* siRNA modified, based on alignment to known human gene sequences. Multiple siRNAs targeting *NDUFS2* and *NDUFS7* were screened to determine the most effective siRNA (**Figure S1A-C**). The sequences for the siRNA used are shown in **Table S1**. The optimal time-point to measure endpoints after siRNA administration was experimentally determined based on depression of protein expression, assessed by immunoblot, measured 24-, 48-, and 72-hours post-siRNA administration (**Figure S1D-G**). Maximal knockdown was achieved at 48-hours in hDASMCs treated with siNDUFS1, siNDUFS2, and siUQCRFS1 (**Figure S1D-F**). In siCOX4I2-treated cells, there was a 38-58% knockdown at 48-hours of treatment, with a further reduction to 13-23% knockdown at 72-hours (**Figure S1G**). For consistency, all endpoint experiments were conducted at 48-hours of treatment.

### 3.2. siRNA Treatment Significantly Knocked Down ETC Subunit mRNA and Protein Expression

All siRNA constructs significantly reduced their target at both mRNA and protein levels. siNDUFS1 significantly decreased the expression of *NDUFS1* mRNA (**Figure 2A**, n=4 cell lines, 93.5% decrease, p=0.0007) and protein (**Figure 2B**, n=7 cell lines, 58.4% decrease, p=0.0034). siNDUFS2 significantly reduced the expression of *NDUFS2* mRNA (**Figure 2C**, n=6 cell lines, 94.3% decrease, p<0.0001) and protein (**Figure 2D**, n=7 cell lines, 87.8% decrease, p=0.0006). siNDUFS7 significantly reduced the expression of *NDUFS7* mRNA (**Figure 2E**, n=6 cell lines, 98.2% decrease, p<0.0001) and protein (**Figure 2F**, n=6 cell lines, 75.6% decrease, p=0.0004). siUQCRFS1 significantly reduced the expression of *UQCRFS1* mRNA (**Figure 2G**, n=6 cell lines, 92.8% decrease, p=0.0014) and protein (**Figure 2H**, n=7 cell lines, 93.9% decrease, p<0.0001). siCOX4I2 significantly reduced the expression of *COX4I2* mRNA (**Figure 2I**, n=3 cell lines, 90.1% decrease p=0.036) and protein (**Figure 2J**, n=5 cell lines, 51.2% decrease p=0.0077).

**Figure 2:**
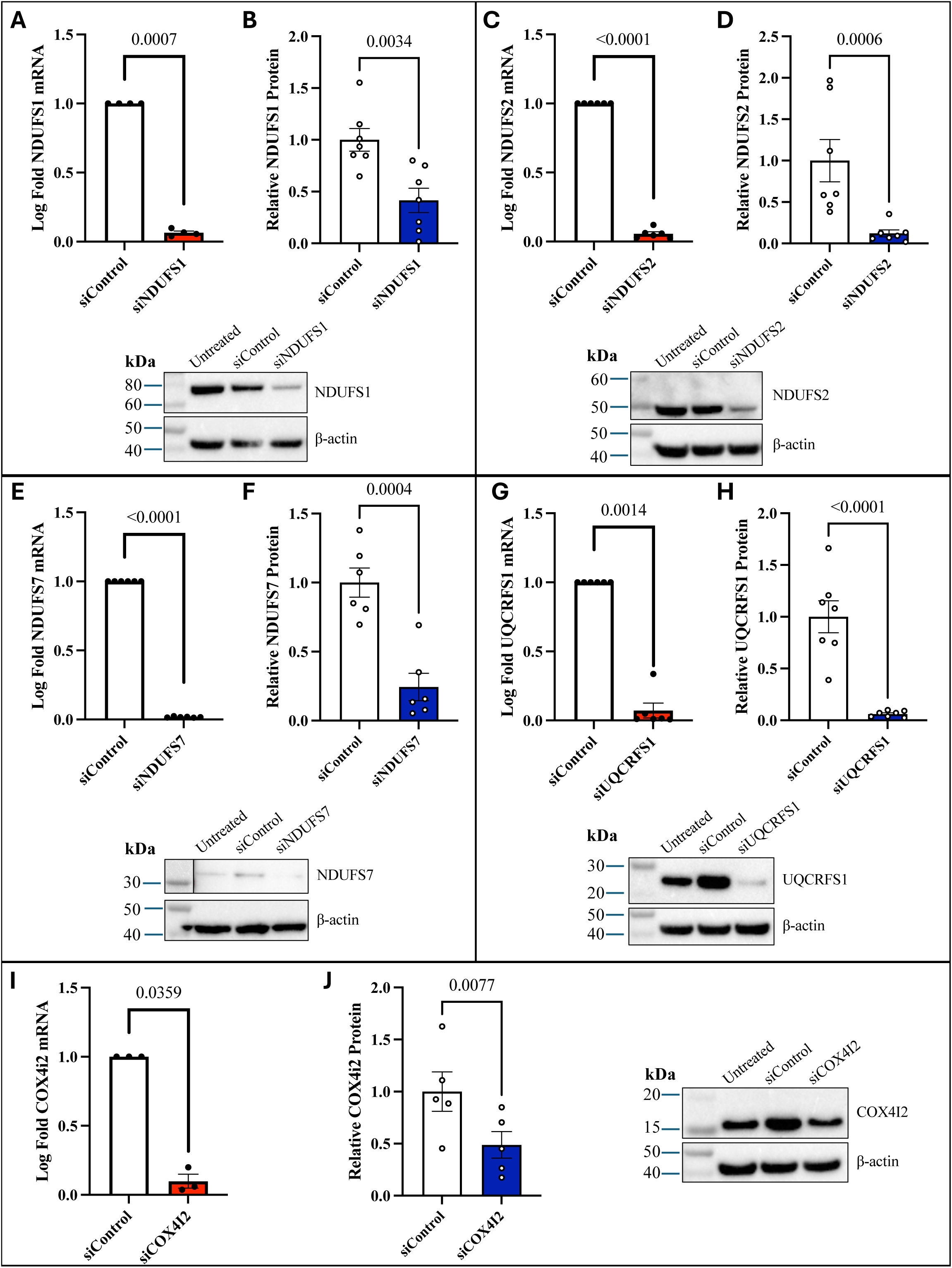
Confirmation of ETC Subunit Knockdown at the mRNA and Protein Level. **A-B**) NDUFS1 mRNA (n=4 cell lines) and protein (n=7 cell lines) were significantly reduced following siNDUFS1 versus siControl. **C-D**) siNDUFS2 significantly decreased expression of NDUFS2 mRNA (n=6 cell lines) and protein (n=7 cell lines) versus siControl. **E-F**) siNDUFS7 significantly decreased expression of NDUFS7 mRNA (n=6 cell lines) and protein (n=6 cell lines) versus siControl. **G-H**) siUQCRFS1 significantly decreased expression of UQCRFS1 mRNA (n=6 cell lines) and protein (n=7 cell lines) versus siControl. **I-J**) siCOX4I2 significantly decreased expression of COX4I2 mRNA (n=3 cell lines) and protein (n=5 cell lines). All measurements made 48-hours post siRNA treatment. Three technical replicates were run for qRT-PCRs. For all immunoblots, representative gels shown and expression normalized to β-actin.

### 3.3. Knockdown of NDUFS2 Significantly Inhibits Oxygen Responsiveness of hDASMCs

Across repeated experiments, we confirmed that the primary hDASMCs retained their unique oxygen responsive phenotype, exhibiting cell shortening upon exposure to normoxic media as assessed by confocal imaging. The mitochondrial network was quantified with the mitochondrial potentiometric dye TMRM (tetramethylrhodamine methyl ester) and mitochondrial fragmentation was objectively quantified using an established machine learning algorithm which classified mitochondria based on length (**Figure S2C-D**). Oxygen increased mitochondrial fission within 10 minutes of the transition from hypoxia to normoxia (**Figure 3A-B**).

**Figure 3:**
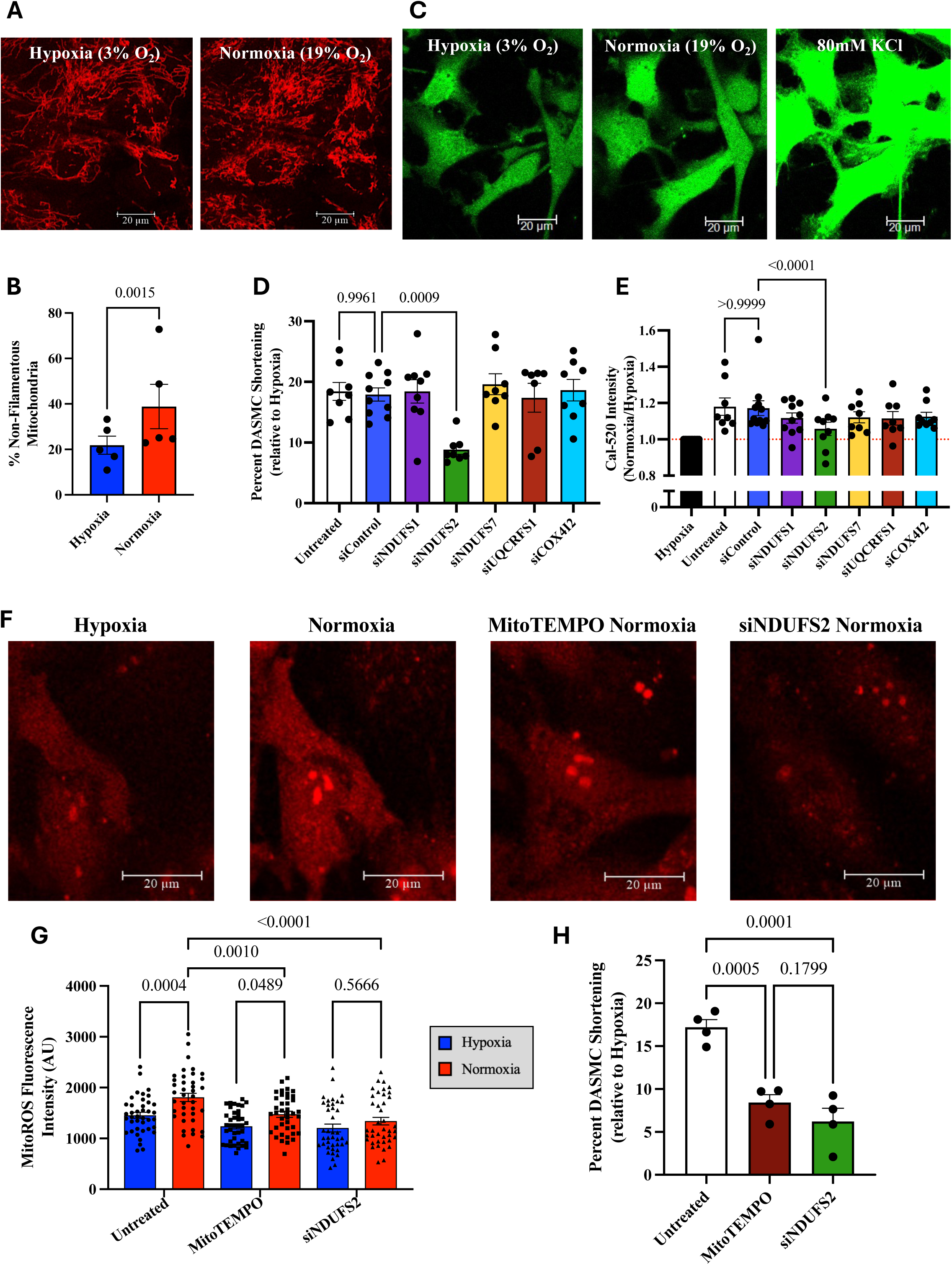
NDUFS2 Mediates ROS Production and Vasoconstriction in Response to Oxygen in human DASMCs. Oxygen responsiveness was assessed using live confocal imaging with the mitochondrial dye tetramethylrhodamine methyl ester (TMRM) and the calcium sensitive dye Cal-520AM. **A**) Representative images of DASMCs stained with TMRM in hypoxia (left, 3% oxygen) and following 20 minutes normoxia (right, 19% oxygen). **B**) Normoxia significantly increased mitochondrial fission, quantified as an increase in the percentage non-filamentous mitochondria (punctate and intermediate mitochondria) (n=5 cell lines). **C**) Representative images of DASMCs stained with Cal-520AM measured after 20-minutes in hypoxia (left, 3% oxygen), 20-minutes normoxia (middle, 19% oxygen), and with the addition of 80mM KCl (right) as a positive control. Cells lacking a KCl-induced rise in intracellular calcium were excluded from analysis. **D**) Treatment with siControl caused no change in normoxia-induced rise in [Ca^2+^]_i_ versus untreated cells, whereas siNDUFS2 significantly reduced the oxygen-induced increase in [Ca^2+^]_i_ (n=10 cell lines, 10-20 cells/cell line). **E**) There was also no effect of siControl on DASMC constriction compared to untreated cells, measured as % cell shortening versus hypoxic baseline, whereas siNDUFS2 significantly decrease normoxia-induced constriction (n=10 cell lines, 10 cells/cell line). **F**) Representative images showing mROS measured using MitoROS™ 580 in hypoxia (3% oxygen) and normoxia (19% oxygen) in untreated cells (left and middle-left), with antioxidant (MitoTEMPO, middle-right), and following NDUFS2 knockdown (right). **G**) In untreated cells, there was a significant rise in mROS generation in normoxia, with a significant decrease in normoxic mROS generation with MitoTEMPO and siNDUFS2 (n=4 cell lines, 10 cells/cell line). **H**) MitoTEMPO significantly decreased oxygen-induced cell shortening, to a similar degree as NDUFS2 knockdown (n=4 cell lines, 10 cells/cell line).

Oxygen responsiveness was further assessed via confocal imaging with the calcium-sensitive dye Cal-520AM, defined as a rise in [Ca^2+^]_i_ within 20 minutes of oxygen exposure (**Figure 3C left and middle**). Challenge with 80mM KCl served as a positive control, stimulating the maximal possible rise in intracellular calcium (**Figure 3C right**). KCl-induced increases in intracellular calcium were unaffected by any siRNA treatment (**Figure S2A** n=10 cell lines, 10-20 cells per cell line). Across n=10 DASMC lines, normoxia increased [Ca^2+^]_i_ 18.6±2.3% in untreated cells (**Figure 3D**), with no significant difference between untreated and siControl cells (16.0±1.9%, p>0.9999). In contrast, siNDUFS2-treated cells had only a 5.5±1.5% increase in [Ca^2+^]_i_ (p<0.0001), versus hypoxic baseline. The knockdown of NDUFS2 had no effect on KCl-induced increases in [Ca^2+^]_i_ (**Figure S2A**), indicating that siNDUFS2 specifically inhibited the oxygen response of the DASMCs. There was a concomitant reduction of oxygen-induced DASMC shortening with siNDUFS2 (**Figure 3E**). Upon oxygen exposure, DASMC length decreased by 18.4±1.1%, relative to the length in hypoxia (n=10 cell lines); siNDUFS2 significantly attenuated this oxygen-induced cell shortening (**Figure 3E**, 8.9±0.7%, p=0.0009).

### 3.4. Oxygen-Induced Generation of Mitochondrial ROS is Inhibited by siNDUFS2

Oxygen-induced generation of mitochondrial superoxide was assessed with live cell confocal imaging using the dye MitoROS™ 580 (n=4 cell lines, 10 cells per cell line) to examine this early step in the DASMC oxygen response pathway (**Figure 1B**). There was a 24±4.9% increase in mitoROS fluorescence intensity upon exposure of untreated cells to normoxia (**Figure 3F-G**, 1456±57.6 to 1807±80.95 AU, p=0.0004). Pre-treatment with a mitochondrially-targeted antioxidant (MitoTEMPO) reduced normoxia-induced mROS (**Figure 3F-G**, to 1475±56.63 AU, p=0.001), as did NDUFS2 knockdown (**Figure 3F-G**, to 1340±75.96 AU, p<0.0001). While MitoTEMPO significantly reduced mROS, a residual increase in response to oxygenation was still detected upon reoxygenation (**Figure 3G**, 1238±49.39 to 1475±58.63 AU, p=0.049). Although statistically significant, this residual rise in mROS was markedly attenuated and did not exceed baseline mROS levels observed in untreated hypoxic cells.

siNDUFS2 also reduced O_2_-induced DASMC shortening (**Figure 3H**, from 17.19±0.91% to 6.22±1.5% shortening, p=0.0001). Likewise, inhibiting mROS generation with MitoTEMPO attenuated oxygen-induced DASMC shortening (**Figure 3H**, from 17.19±0.91% to 8.43±0.91%, p=0.0005).

### 3.5. Knockdown of ETC Subunits, with the Exception of NDUFS1, Did Not Alter Complex I, III, and IV Activity at 48 hours

Activity of Complex I was measured by oxidation of NADH by dipstick-immunocaptured Complex I (**Figure 4A**). Complex I activity of hDASMCs (n=8 cell lines) transfected with siRNA targeting NDUFS2, NDUFS7, UQCRFS1, and COX4I2 were not significantly different than siControl-transfected cells (**Figure 4B**). Knockdown of NDUFS1 significantly reduced Complex I activity, reported as signal intensity in arbitrary units (AU): siControl=2287±250, siNDUFS1=1775±199, p=0.024. The assay measured activity as NADH oxidation of immunocaptured Complex I, and with the localization of NDUFS1 close to the NADH binding site, the observed slight reduction in Complex I activity following NDUFS1 knockdown was anticipated.

**Figure 4:**
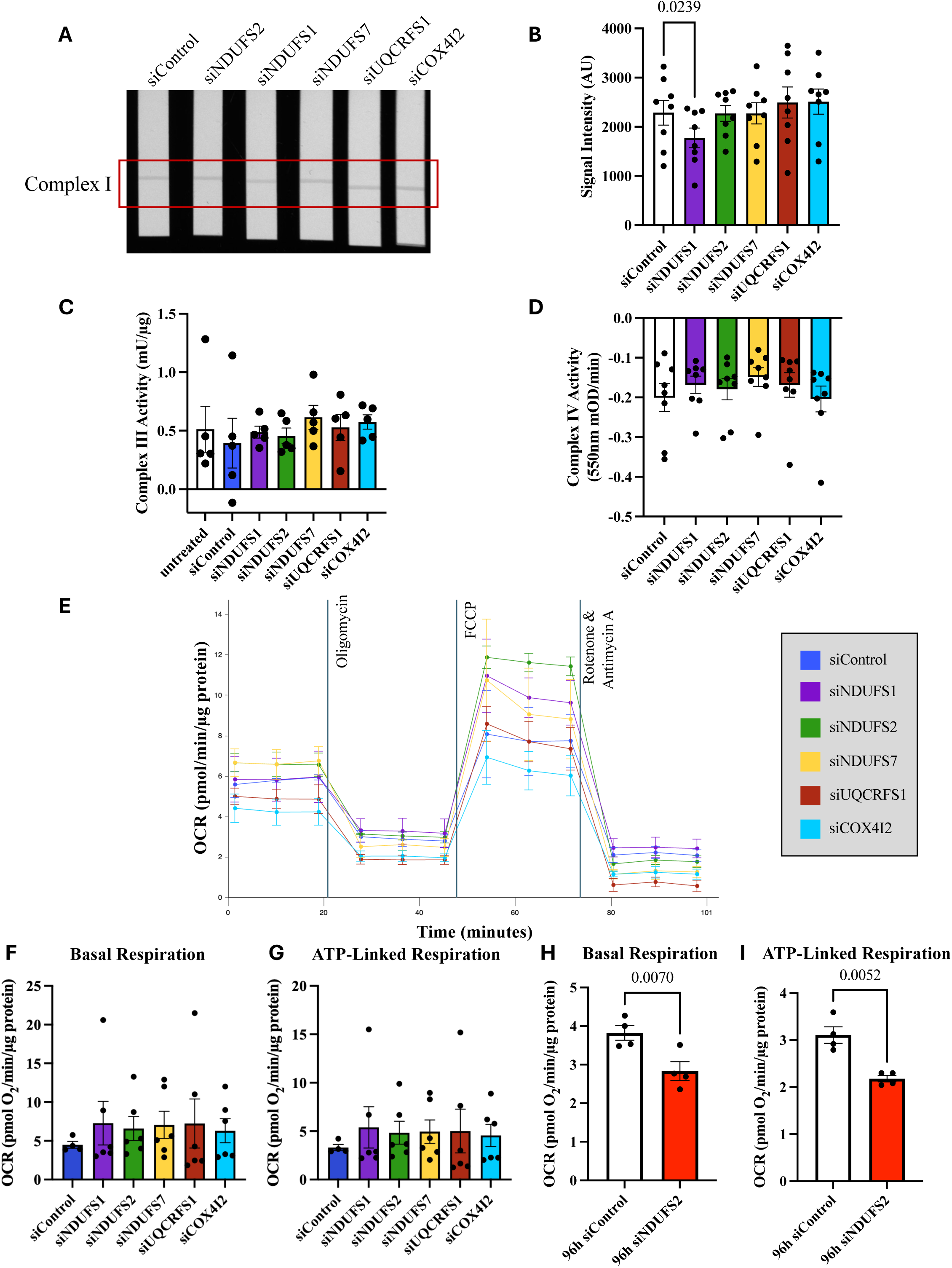
Selective Knockdown of ETC Subunits Does Not Inhibit Mitochondrial Subunit Function or Respiration. **A**) Complex I activity was measured using a dipstick assay of hDASMCs treated for 48h with siRNA targeting control, NDUFS2, NDUFS1, NDUFS7, UQCRFS1, and COX4I2. **B**) Complex I activity was only significantly reduced by treatment with siNDUFS1 (n=8 cell lines). **C**) Complex III activity, normalized to µg of protein, was not significantly changed by treatment of any siRNA including siUQCRFS1 (n=5 cell lines). **D**) There was no significant difference in Complex IV activity with 48h treatment with any siRNA treatment, including siCOX4I2 (n=8 cell lines). **E**) A Seahorse XF mitochondrial stress test measured oxygen consumption rate (OCR) of hDASMCs treated for 48-hours with siRNA targeting control (NC) or ETC subunits: NDUFS2, NDUFS1, NDUFS7, UQCRFS1, or COX4I2. Knockdown of ETC subunits (n=5 cell lines, 3 technical replicates) caused no change in key measures of mitochondrial respiration, including (**F**) basal respiration or (**G**) ATP-linked respiration. **H**) 96-hours treatment of hDASMCs with siNDUFS2 significantly decreased basal respiration and **I**) ATP-linked respiration as measured with a Seahorse XF mitochondrial stress test. OCR in each cell line and treatment was normalized to total µg protein.

Complex III activity was measured as the change in concentration of reduced cytochrome c over time, measuring a linear subset within a recorded 10-minute period (**Figure S3A**). Complex III activity was calculated based on a standard curve (not shown), subtracting the change in concentration of reduced cytochrome c, over time, in samples treated with Antimycin A (**Figure S3B**). Activity of Complex III in hDASMCs (n=5 cell lines) transfected with siRNA targeting NDUFS1, NDUFS2, NDUFS7, UQCRFS1, COX4I2, or siControl was not significantly different (p=0.38) from activity in cell lines grown for 48-hours without siRNA (**Figure 4C**).

The activity of Complex IV was determined colorimetrically, measuring the change in absorbance at 550 nm to follow the oxidation of reduced cytochrome c and subtracting the background absorbance slope (**Figure S3C**). Across n=8 hDASMC cell lines, there was no significant change (p=0.58) in Complex IV activity with knockdown of NDUFS1, NDUFS2, NDUFS7, UQCRFS1, or COX4I2, compared to siControl transfected cells (**Figure 4D**), indicating that ETC subunit knockdown did not impair the overall function of Complexes I, III, or IV.

### 3.6. ETC Subunit Knockdown Did Not Alter Mitochondrial Metabolic Function at 48 hours

Whole cell mitochondrial metabolism was assessed with micropolarimetry using the Seahorse Mito Stress Test (**Figure S4A**), using an experimentally optimized concentration of FCCP (**Figure S4B**). Transfection of hDASMCs (n=6 cell lines) with siRNA for 48-hours did not significantly change key mitochondrial metabolic parameters (**Figure 4E**), including basal respiration (**Figure 4F**) and ATP-linked respiration (**Figure 4G**). There was similarly no effect of any ETC subunit knockdown on other derived mitochondrial metabolic parameters (**Figure S4C-H**). At this 48-hour timepoint of functional assessments of the effects of ETC subunit knockdown on oxygen responsiveness, there was no suppression of mitochondrial metabolic function observed. However, due to the integral role of NDUFS2 in Complex I assembly and function it would be expected that ultimately a decrease in NDUFS2 should reduce respiration. Consequently, we assessed the effects of NDUFS2 knockdown on mitochondrial function 96-hours post-siRNA transfection using the Seahorse Mito Stress Test. Following 96-hours of siNDUFS2 transfection, there was indeed a suppression in basal respiration (**Figure 4H**, n=4 cell lines, p=0.007) and ATP-linked respiration (**Figure 4I**, n=4 cell lines, p=0.0052), with no effect of 96-hours of siNDUFS2 transfection on other derived mitochondrial metabolic parameters (**Figure S5A-G**).

### 3.7. Knockdown of ETC Subunits Induced Widespread Transcriptomic Changes, with NDUFS2 Knockdown inducing a Robust Change in Mitochondrial Transcriptome

Treatment of hDASMCs (n=5 cell lines) with siRNA targeting NDUFS2, NDUFS1, NDUFS7, UQCRFS1, or COX4I2 for 48-hours induced substantial transcriptomic changes compared to siControl, as assessed via 3’RNA sequencing. Comprehensive enrichment analyses and complete gene lists are available in the supplement, with raw and processed data available through NCBI’s gene expression omnibus (GSE335748). There were 677 differentially expressed genes (DEGs) commonly regulated between siControl versus ETC knockdowns that were found independent of which subunit was knocked down (**Figure 5A, Figure S6**).

**Figure 5:**
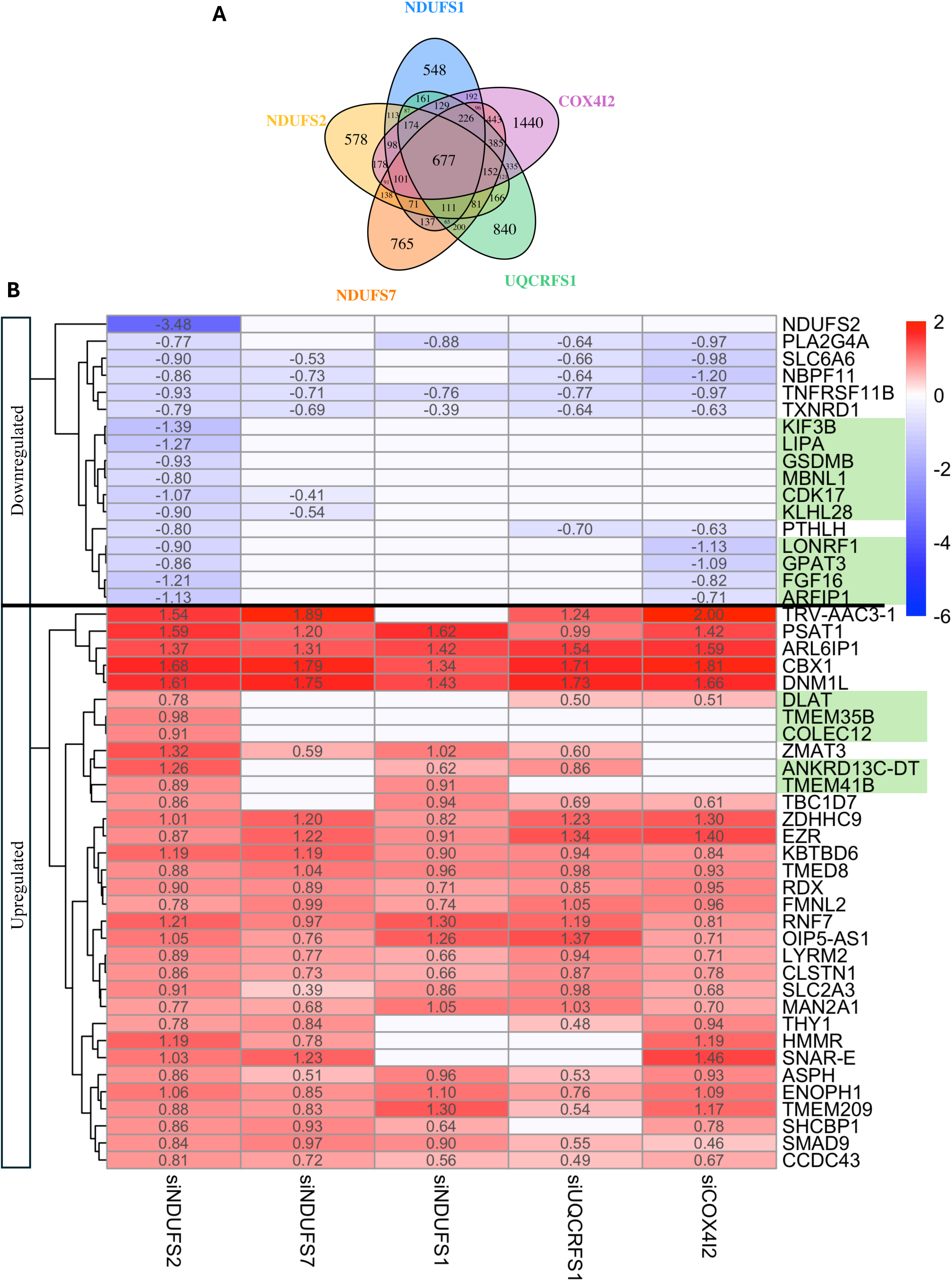
Transcriptomics Reveal Unique Downstream Effects of Selective Knockdown of ETC Subunits on Gene Expression. **A**) There were numerous differentially expressed genes (DEGs) significantly regulated by each knockdown condition compared to siControl, with 677 common DEGs across all knockdown conditions. **B**) Heat map of the top 50 siNDUFS2 versus siControl DEGs with each siRNA, reveals both unique (highlighted in green) and common regulation of genes amongst knockdown conditions. Colour was scaled by fold change achieved by each knockdown versus siControl.

To compare the effects of siNDUFS2 to other ETC knockdowns, we examined the log-fold change of each knockdown versus siControl for the top 20 DEGs between siNDUFS2 and siControl (**Figure 5B**). siNDUFS2 exhibited a distinct gene ontology (GO) phenotype among knockdown conditions, uniquely downregulating *NDUFS2*, *FGF16*, *AFRIP1*, *CDK17*, *KIF3B*, *GSDMB*, *KLHL28*, *LONRF1*, *GPAT3*, and *LIPA*, and uniquely upregulating *DLAT*, *TMEM35B*, *COLEC12*, *TMEM41B*, and *ANKRD13C-DT*, as discussed below (**Figure 5B**). Of these 20 DEGs, there was also a common effect of ETC subunit knockdowns on expression of genes, such as the histone-binding *CBX1* and *DNM1L*, the gene encoding DRP1, the major mediator of mitochondrial fission. There was also a shared effect of some knockdown conditions on expression of the mitochondrial fission mediator *DNM1L*, the phosphoserine aminotransferase *PSAT1*, the epigenetic regulator *CBX1*, and the valine tRNA *TRV-AAC3-1* **(Figure 5B**).

Knockdown of NDUFS2 uniquely altered the expression of 578 genes relative to siControl (**Figure 5A**), meaning these changes were not observed following siRNA-mediated silencing of other ETC subunits. GO analysis of these unique 578 genes revealed enrichment of numerous pathways related to mitochondrial structure and functions in the top 20 GO pathways (**Figure 6A-C**), with most of the genes captured within these mitochondrial GO pathways being upregulated by NDUFS2 knockdown (**Figure 6D**).

**Figure 6:**
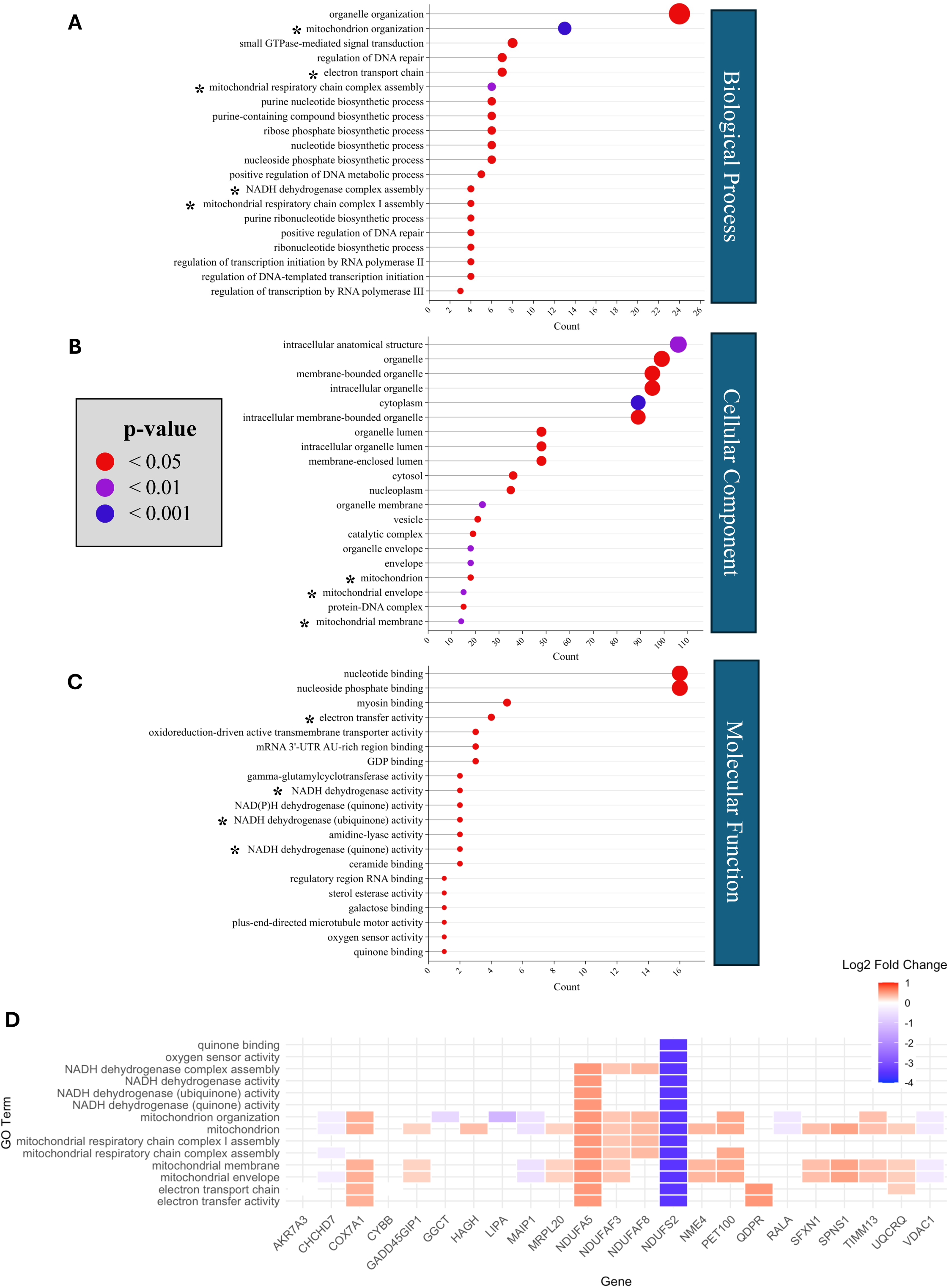
Gene Ontology (GO) Enrichment Analysis of Differentially Expressed Genes (DEGs) Uniquely Regulated by NDUFS2 Knockdown Reveals Key Mitochondrial Pathways. The top 20 significantly enriched (adjusted p<0.05) GO terms in the (**A**) Biological Process, (**B**) Cellular Component, and **(C**) Molecular Function domains, ranked by the number of genes involved in each GO term (count). Mitochondrially related pathways are indicated by an asterisk (*). **D**) The heatmap of DEGs within the mitochondrially related pathways (* in **A-C**) demonstrates the majority of the mitochondrial genes uniquely regulated by siNDUFS2, among the 5 knockdown conditions, are upregulated by siNDUFS2 compared to siControl. Colour was scaled by log2 fold change of siNDUFS2 versus siControl.

### 3.8. Unique Transcriptomic Changes Induced by NDUFS2 Knockdown Among ETC Complex I Subunit Knockdowns

To determine the effects of NDUFS2 knockdown within the I_Q_ site, the significantly DEGs between siControl and siNDUFS2 were compared to significantly DEGs with siNDUFS7 versus siControl (**Figure 7A**). The top 20 enriched GO pathways for the 1,515 genes regulated by siNDUFS2 but not siNDUFS7 revealed many mitochondrial pathways, with most of the genes contained in these pathways being upregulated by siNDUFS2 (**Figure 7B & 7D**). The top 20 enriched GO pathways of the 2,317 genes regulated by NDUFS7 but not siNDUFS2 did not include mitochondrial pathways (**Figure S13B**), nor did the top 20 pathways enriched of the 1,422 genes commonly regulated by siNDUFS2 and siNDUFS7, with the commonly regulated genes enriched for pathways relating to growth and development (**Figure S13C-D**).

**Figure 7:**
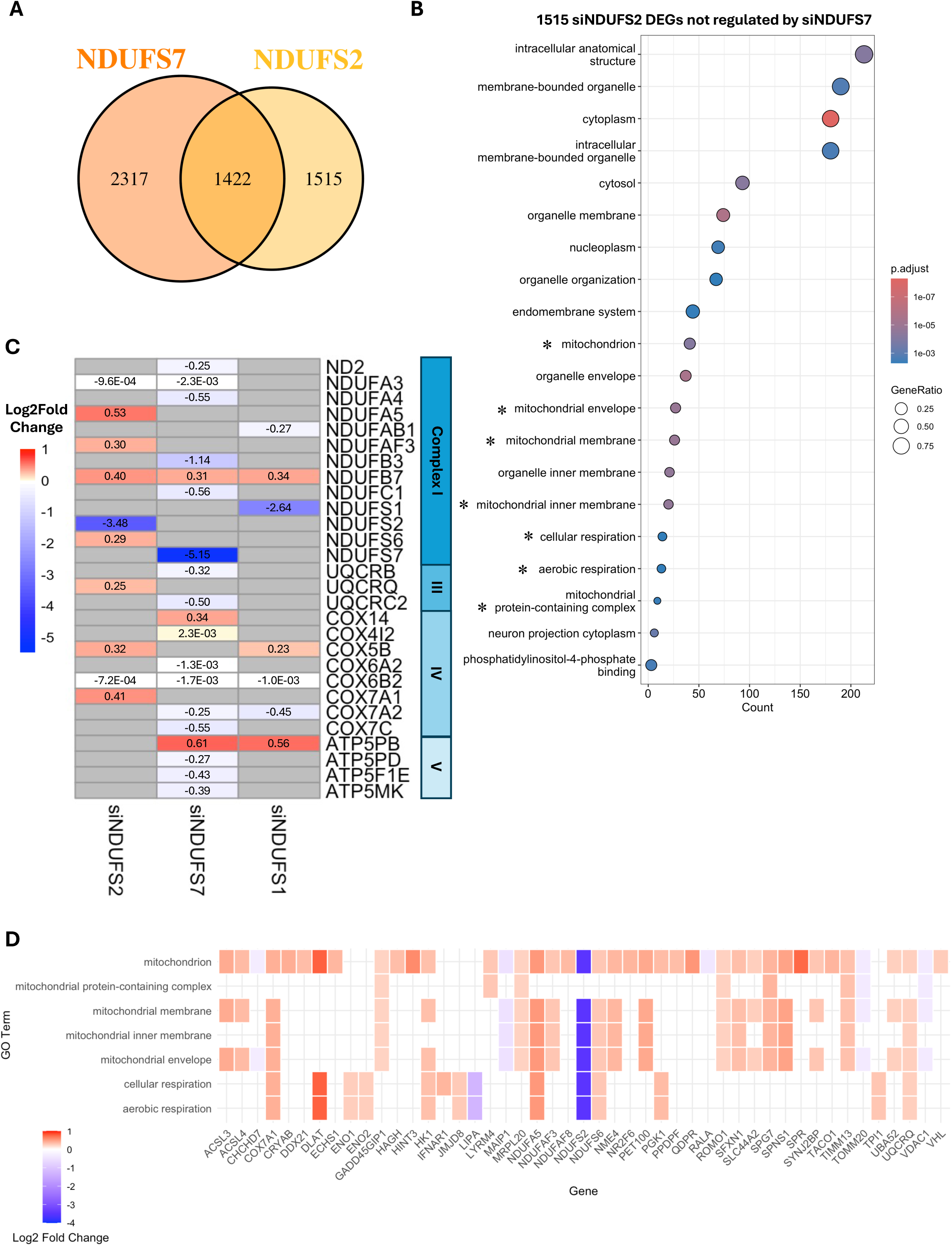
Unique Mitochondrial Transcriptomic Effect of NDUFS2 Knockdown Among ETC Complex I Knockdowns. **A**) Comparison of significant (adjusted p<0.05) DEGs between siControl and siNDUFS7 (left) and between siControl and siNDUFS2 (right) to examine the differential effects of knocking down neighboring subunits in the ubiquinone binding site of ETC Complex I. **B**) The top 20 significantly enriched (adjusted p<0.05) GO terms of the 1515 genes regulated by NDUFS2 knockdown but not NDUFS7 knockdown, selecting the most significantly enriched pathways (smallest p-value) then arranged by the number of DEGs within each pathway (count). Mitochondrially related pathways are indicated by asterisks (*). **C**) Comparison of significantly differentially expressed (adjusted p<0.05) ETC components between knockdowns of Complex I subunits (siNDUFS2, siNDUFS7, and siNDUFS1). siNDUFS2 induced a potentially compensatory upregulation of more ETC components than either other Complex I knockdown, with the largest effect in Complex I. NDUFS1 knockdown affected the fewest other ETC genes while NDUFS7 knockdown downregulated many ETC genes. **D**) The heatmap of DEGs within the mitochondrially related pathways (* in **B**) demonstrates the majority of the mitochondrially genes regulated by siNDUFS2, but not siNDUFS7, are upregulated by siNDUFS2 compared to siControl. Colour was scaled by log2 fold change of siNDUFS2 versus siControl.

In a comparison of the significantly differentially expressed ETC subunit genes across Complex I knockdowns (each compared to siControl), siNDUFS2 upregulated more ETC components than siNDUFS7 or siNDUFS1, with siNDUFS2 having the largest effect on other Complex I genes (**Figure 7C**).

Comparing the top 50 DEGs with NDUFS7 knockdown (**Figure S12**) or NDUFS1 knockdown (**Figure S15**) across all knockdown conditions, there is minimal overlap of the top 50 DEGs from either Complex I knockdown and siNDUFS2 significantly DEGs, with the top DEGs with siNDUFS7 or siNDUFS1 having unique expression profiles among knockdown conditions. A comparison of DEGs with NDUFS2 knockdown and NDUFS1 knockdown (**Figure S16A**) also demonstrated enrichment of mitochondrial and metabolic pathways for the 1,506 gene regulated by siNDUFS2 but not siNDUFS1, with most of the genes within the mitochondrial pathways being upregulated by siNDUFS2 (**Figure S16C-D**). A similar enrichment of mitochondrial pathways was not seen with the 1,554 genes regulated by siNDUFS1 but not siNDUFS2 (**Figure S16B**), nor in the 1,431 genes commonly regulated by siNDUFS1 and siNDUFS2 (**Figure S16E**). Examining all enriched mitochondrial GO pathways for the DEGs uniquely and commonly regulated by siNDUFS7 and siNDUFS2 (**Figure S14**) and for the DEGs uniquely and commonly regulated by siNDUFS1 and siNDUFS2 (**Figure S17**), most of the mitochondrial genes regulated by siNDUFS2 were consistently upregulated by siNDUFS2, while the mitochondrial genes uniquely regulated by siNDUFS7 or siNDUFS1 were mostly downregulated by the Complex I knockdowns.

## 4. Discussion

We previously identified downregulation of NDUFS2 mRNA in hDASMCs in response to sustained normoxia^28,29^. Here we examined the functional contribution of candidate ETC subunits to oxygen-induced hDASMC constriction, focusing on NDUFS2 and other putative oxygen-sensor subunits described in HOSS tissues. We assessed the functional effects of ETC subunit knockdown, while also performing an RNAseq analysis to establish the transcriptomic impact of selectively changing the expression of each of these genes.

We demonstrate for the first time that NDUFS2 mediates the human DASMC’s oxygen response. Only knockdown of NDUFS2 inhibited the oxygen-induced rise in intracellular calcium and decrease in cell length, surrogates for DA constriction *in vivo.* siNDUFS2 also significantly attenuated the oxygen-induced rise in mitochondrial superoxide in hDASMCs, indicating that it likely inhibited DASMC contraction by interfering with the mROS signaling mechanism. Consistent with this, inhibiting mROS also inhibited oxygen-induced constriction of DASMC. It is noteworthy that siNDUFS2 knockdown had no effect on KCl-induced rise in intracellular calcium, supporting the specificity of its effects for the response to oxygen. The loss of DA oxygen responsiveness induced by NDUFS2 knockdown occurred independently from effects on mitochondrial metabolic function, with no detectable changes in mitochondrial respiration or ETC Complex I, III, or IV activity at the same time point (48 hours) at which oxygen sensing was inhibited. Thus, NDUFS2 plays a vital, early role in oxygen-induced DA constriction through its ability to reduce the physiologic increase in mROS that occurs in response to increases in pO_2_.

Importantly, we also demonstrated that these DAMSC, each isolated from an individual infant’s DA at the time of congenital heart surgery, were indeed SMC and that, despite passage in hypoxic culture, they retained the unique, classical DA oxygen responsive phenotype, manifesting as an increase in mitochondrial fission^13^, rise in cytosolic calcium, and contraction in response to physiologic oxygen concentrations, as occurs with the first breath^28^.

NDUFS2 is highly conserved and is an essential component on Complex I^33^, yet 48-hours following transfection with siRNA we did not observe a decrease in mitochondrial respiration or ETC Complex I, III, or IV activity, although by 96-hours basal oxygen consumption was depressed. Our results indicate a temporal disconnect between the early effects of NDUFS2 knockdown (inhibiting hDASMC acute oxygen responsiveness, **Figure 3,** 48-hours) versus the later effects (depression of basal and ATP-linked mitochondrial respiration, **Figure 4H-I**, 96-hours). This suggests short-term compensation of mitochondrial respiration (**Figure 4E-G**), potentially via Complex II, while mitochondrial generation of superoxide is suppressed. However, as NDUFS2 is part of the minimal assembly of Complex I^33^, one might expect that reducing the expression of NDUFS2 would ultimately reduce the cell’s bioenergetic status, and indeed it does at 96-hours. However, in the initial 48-hours after administration of siNDUFS2, electron leak and ROS production, which are key to oxygen sensing and signal transduction in DASMCs, appear more sensitive to siNDUFS2 than does ATP production. This is consistent with the findings of Brand et al., who developed small molecule electron leak suppressors, like S1QEL and S3QEL^34,35^. S1QEL inhibits electron leak and formation of superoxide at the I_Q_ site in ETC Complex I, without simultaneously reducing ATP^34,36^. We previously showed that S1QEL inhibits oxygen sensing and DA constriction in rabbit DAs, without impairing phenylephrine-induced DA constriction, ETC Complex I activity, or bioenergetics^25^. This suggests the production of signalling ROS, though a byproduct of electron flux, operates in parallel or at different sensitivity to bulk electron flux through Complex I. We also suggest an additional mechanism by which metabolic flux may be conserved, at least temporarily, following knockdown of NDUFS2. Transcriptomic results show siNDUFS2 uniquely upregulated many components of ETC Complex I, including NDUFA5, NDUFAF3, NDUFB7, and NDUFS6 (**Figure 7C**), as well as upregulating many other mitochondrial genes found in related GO pathways (**Figures 6D, 7D, S14D, S16D, and S17D**). This suggests a compensatory, transcriptional remodelling of ETC Complex I, which may explain the temporary preservation of bioenergetics, which we documented by micropolarimetry 48-hours after exposure to siNDUFS2. This would align with the findings of Bandara et al., who demonstrated the essential role of NDUFS2 in ROS generation and found a compensatory increase in Complex II respiration following CRISPR/Cas9-mediated mutation of NDUFS2 in HEK293 cells^37^.

Knockdown of other Complex I subunits, NDUFS1 and NDUFS7, did not affect oxygen responsiveness or mitochondrial respiration at the 48-hour time point. Although, knockdown of NDUFS1 decreased proximal Complex I activity (i.e. NADH oxidation). NDUFS2, NDUFS1, and NDUFS7 are all highly conserved subunits, part of the 14 subunits of minimal Complex I assembly^33^. NDUFS2 and NDUFS7 are situated at the ubiquinone binding site (I_Q_), where electrons are transferred to ubiquinone and where rotenone, the classical Complex I inhibitor, binds^38^. This I_Q_ site is a major site of mROS generation in Complex I^39,40^; mROS generation from this site mediates rabbit DA closure *in vivo*^25^. The absence of effect of NDUFS7 knockdown on DA oxygen response indicates a unique role for NDUFS2 within the I_Q_ site in oxygen sensing, likely through its ability to alter mROS generation.

NDUFS1, in contrast, is in the peripheral arm of Complex I (**Figure 1**), part of the electron input/dehydrogenase (N) functional module of Complex I that accepts electrons via oxidation of NADH. While it is the prosthetic group flavin mononucleotide (FMN), and not NDUFS1 that directly accepts electrons from NADH^41^, NDUFS1 plays a vital role in the function and stability of the N-module^42^. NDUFS1 is associated with multiple iron-sulfur (Fe-S) clusters in Complex I that are essential cofactors and mediate electron transfer in the respiratory chain, including Fe-S clusters N1b, N4, and N5^38,43^. A recent study demonstrated that biallelic mutation of NDUFS1 decreased the N-module stability and disrupted the flow of electrons between two Fe-S clusters (N4 and N5), impairing supercomplex formation in patient-derived skin fibroblasts^42^. With the contribution of NDUFS1 to the N-module structure and function, the inhibition of Complex I function that we observed with knockdown of NDUFS1 was expected, particularly as the assay measured activity based on NADH oxidation. However, the lack of effect of NDUFS1 knockdown on live cell mitochondrial respiration suggests respiratory chain function was maintained at 48-hours, despite this knockdown. One might expect bioenergetic impairment at a later timepoint following siNDUFS1 knockdown, similar to that observed 96-hours following siNDUFS2 treatment; however, this study lacked the timepoint granularity to capture if or when there was no longer bioenergetic compensation following ETC subunit knockdown.

It is well known that mROS in DASMCs initiate the DA response to oxygen^14^. Complex I and III are the primary sites of mROS generation within the mitochondria^24^, with the diffusible ROS (e.g. H_2_O_2_) serving as a redox messenger to alter the activity of Kv channels and thus the polarization/depolarization state of the DASMC membrane^21^. Similar to the effects seen in the adult PA^23^, treatment of DASMCs with classic inhibitors of both ETC Complex I (rotenone) and Complex III (antimycin A) mimics hypoxia^14^. However, these inhibitors not only stop the flow of electrons through the ETC thereby potentially eliminating mROS signaling molecules; they also suppress metabolism making it difficult to differentiate their effects on Complex I or III mROS generation from their bioenergetic and ETC inhibitory effects. Thus, we previously used a more specific pharmacologic probe, S1QEL (suppressor of site I_Q_ electron leak)^34^ to inhibit Complex I ROS generation without altering bioenergetics. S1QEL inhibited oxygen-induced rabbit DA constriction *ex vivo* and prevented rabbit DA closure at birth *in vivo*^25^. This response was unique to Complex I mROS, with no effect of inhibiting Complex III electron leak using S3QEL (suppressor of site III_Qo_ electron leak)^35^ on oxygen-induced rabbit DA constriction *ex vivo*, with preserved vasoconstriction to KCl and phenylephrine following both S1QEL and S3QEL treatment^25^. These early studies identify electron leak and ROS, rather than metabolism, as the sensor mechanism with Complex I. The current study advances that finding by identifying NDUFS2, situated in the I_Q_ site of Complex I, as the primary source of oxygen-induced mROS generation in humans and demonstrating its key role in oxygen-induced DASMC constriction (**Figure 3F-H**).

Further supporting this conclusion, inhibition of mROS using the mitochondrial superoxide scavenger MitoTEMPO, at a concentration that decreased normoxic mROS levels, attenuates oxygen-induced cell shortening to a similar degree as NDUFS2 knockdown (**Figure 3H**). Together these results demonstrate that knockdown of NDUFS2 inhibits DASMC oxygen responsiveness via inhibition of mROS generation within ETC Complex I. These results support the paradigm that the DASMC oxygen response results from oxygen-induced increases in mROS from Complex I^14,25,44^.

Complex III mROS generation has not been intensively studied in DA oxygen sensing (although we found no effect of S3QEL on DA constriction in rabbits); however, it is not clear whether Complex I or III is most responsible for initiating hypoxic pulmonary vasoconstriction (HPV)^27,31^. Based on this disagreement in the adult oxygen sensing literature, as well as the known hypoxia mimicking effect of antimycin A^14^, we sought to determine any possible role of Complex III in the DA oxygen response by knocking down the Rieske Fe-S cluster. Successful knockdown of UQCRFS1 (confirmed at the protein level) had no effect on oxygen responsiveness, mitochondrial respiration, or ETC Complex activity (similar to our previous observations in adult PASMCs^27^), arguing against an important role for complex III in the human DASMC’s acute oxygen response.

Our examination of Complex IV subunit COX4I2 was similarly based on evidence that it may contribute to oxygen-sensing (HPV) in adult PAs^17^. Complex IV catalyzes the final step of the ETC, receiving the terminal electron from the ETC and converting oxygen to water. COX4I2 modulates Complex IV activity and thus ETC activity^32^. Sommer *et al.* demonstrated impaired HPV in adult COX4I2 knockout mice^32^, though Dunham-Snary *et al.* found no effect of siCOX4I2 on HPV in adult rats^27^. Our incomplete knockdown of COX4I2, was consistent with previously reported levels (∼55% knockdown)^27^; siRNA dose was not increased because complete *COX4I2* knockdown would likely interfere with the function of other ETC Complexes^45,46^. The ETC Complexes I-IV interact and form large supercomplexes^47^, with an interdependence of complex biogenesis and/or stability, particularly that of Complex I^48–50^. Deficiencies in Complex III^48,49^ or IV^45,51^ greatly decrease assembled Complex I in monomeric and supercomplex form. With 51.2±13% knockdown of COX4I2 protein seen in our cells, there was no significant effect on mitochondrial respiration or activity of Complex I, III, or IV. There was also no evidence of COX4I2 contributing to the oxygen response of human DASMCs.

While only NDUFS2 knockdown attenuated DASMC oxygen responsiveness, transcriptomics revealed that all ETC subunit knockdowns regulated DASMC gene expression, with 677 common DEGs between all ETC knockdown conditions versus siControl (**Figure 6A**). GO pathway analysis reveals commonly regulated genes relate to generic cellular processes (**Figure S6A-C**) with no clear mitochondrial phenotype emerging as being common amongst all ETC knockdowns.

The RNA sequencing data also confirms that each knockdown was effective at reducing the expression of the target subunit genes without altering expression of mRNA for the other candidate sensor subunits (**Figure S7A-E**). We were unable to detect *COX4I2* under any siRNA condition in our RNA sequencing dataset (**Figure S7E**); however, we effectively knocked COX4I2 down at both the gene and protein level (as measured by PCR and immunoblot), and there was a clear effect of siCOX4I2 on other gene expression (**Figure 6A)**. 3’ RNA sequencing captures only 75 base pairs from the 3’ end of mRNA, so we hypothesize that seq reads aligned to a shared locus, rendering those reads to be unassigned or assigned to another gene. Others have observed a compensatory upregulation of COX4I2 with deficiency of the other COX4 isoform (COX4I1)^52^, thus is it possible that this interdependent relationship could hold true in the opposite direction; we observed a non-significant increase in *COX4I1* in cells treated with siCOX4I2, further suggesting that COX4I2 was effectively knocked down in our samples.

To compare the effects of the different ETC knockdowns on gene expression, we explored the 50 most differentially expressed genes that resulted from NDUFS2 knockdown across all knockdowns (**Figure 5B**). The genes uniquely upregulated by siNDUFS2 included *DLAT*, *TMEM35B*, *TMEM41B*, *COLEC12*, and the long non-coding RNA (lncRNA), *ANKRD13C-DT*. *DLAT* is a component of the mitochondrial pyruvate dehydrogenase (PDH) complex, playing a key role in mitochondrial metabolism and implicated as a key mediator of granulosa cell proliferation^53^, as well as the tumorigenesis and metabolic reprogramming of multiple cancers^54,55^. *TMEM41B*, an ER-localized transmembrane protein, regulates autophagy and lipid mobilization^56,57^. While there is no known role of these genes in the DA, the unique upregulation of *TMEM41B* and *DLAT* by siNDUFS2 may be indicative of a metabolic shift in these cells with a potential increase in lipid mobilization and PDH activity that warrants further investigation.

*ANKRD13C-DT* (also known as *HHLA3*), which was upregulated by siNDUFS2, has been associated with worse prognosis in multiple solid tumors. Zhou et al. identified *ANKRD13C-DT* as a pyroptosis-related lncRNA in human lung adenocarcinomas, with knockdown decreasing cell proliferation and increasing ROS generation^58^. *TMEM35B*, a transmembrane protein, and *COLEC12*, a pattern recognition molecule of the innate immune system, have also both primarily been studied in cancers and both promote cell proliferation and migration^59–61^. Long non-coding RNAs have been linked to congenital heart diseases and associated lung diseases, including pulmonary arterial hypertension and bronchopulmonary dysplasia^62,63^. However, *ANKRD13C-DT* has not been identified as related to congenital heart diseases nor is it known to have a specific function in the DA, though its effects on ROS generation^58^ may explain its upregulation with NDUFS2 knockdown. Furthermore, the effects of *TMEM35B*, *COLEC12*, and ANKRD13C*-DT* on proliferation may point to a potential role in permanent anatomical DA closure, suggesting a possible multifaceted connection between NDUFS2 manipulation and both functional and anatomical DA closure that merits future study.

Among the top 50 genes regulated by NDUFS2 knockdown, there was unique siNDUFS2-mediated downregulation of ten genes, *ARFIP1, KIF3B, CDK17, FGF16*, *LIPA*, *GSDMB*, *MBNL1*, *KLHL28*, *LONRF1*, and *GPAT3*. *ARFIP1* and *KIF3B* are both involved in intracellular movement and transport, with *ARFIP1* functioning in secretory granule biogenesis and exit from the *trans*-Golgi network^64,65^, and *KIF3B* functioning in mitosis, meiosis, and transport of macromolecules^66^. *CDK17* is an atypical cyclin-dependent kinase^67^ and little is known about its physiological and pathological roles^68^, though it is thought to be involved in the regulation of vesicle internalization and trafficking^68^. *FGF16* is involved in embryonic development and induces cardiomyocyte proliferation^69^, while *MBNL1* is an RNA-binding protein that regulates alternative splicing^70^. *GPAT3* and *LIPA* play key roles in lipid metabolism, with *GPAT3* regulating triglyceride and phospholipid synthesis^71^, and *LIPA* hydrolyzing cholesterol and triglyceride esters in the lysosome^72^. *KLHL28* and *LONRF1* are both involved in protein degradation, with *KLHL28* being a member of the family of substrate adaptors that recognize substrates for ubiquitination^73^, and *LONRF* family proteins containing Lon substrate binding domains for degradation of abnormal and damaged proteins^74^. *GSDMB* is a gasdermin-family pore-forming protein that causes membrane permeabilization and pyroptosis, with isoform specific associations with pro-tumorigenic and pro-metastatic phenotypes in breast cancer^75,76^. These proteins do not have a known role in the pathophysiology of DA closure and oxygen sensing, though another lipase (*LIPC*) was associated with development of PDA^77^ and *CDK17* was identified in a meta-analysis by Yarboro *et al.* as being enriched in the DA versus aorta in rodents^78^. The downregulation of these genes with NDUFS2 knockdown may indicate a change in internalization and secretion of macromolecules, protein degradation, and/or proliferation relating to DA closure, though the specific functions of these genes in the DA remain unknown.

Comparing the top 20 GO pathways in each domain of the DEGs unique to each knockdown condition, the enriched GO pathways for siNDUFS2 had a distinctly mitochondrial phenotype (**Figure 6A-C**). Of the top 20 most enriched GO pathways unique to the other ETC knockdowns (**Figures S8-S11**), the only mitochondrial GO term to appear is “mitochondrion” with siNDUFS1, as compared with the 14 mitochondrial GO terms appearing in the most enriched pathways with siNDUFS2. The genes in mitochondrial and metabolic GO pathways were mostly upregulated by siNDUFS2 (**Figure 6D, 7D, S14D, S16D, S17D**), and among Complex I knockdowns siNDUFS2 upregulated the most ETC subunits, with a stronger effect on Complex I subunits than siNDUFS1 or siNDUFS7. This may indicate a compensatory upregulation of mitochondrial and ETC genes to maintain mitochondrial function with the loss of an integral ETC subunit.

GO analysis of the genes unique to NDUFS2 knockdown also revealed enrichment of nucleotide/nucleoside biosynthetic and binding pathways; these were similarly enriched by chronic normoxic exposure in human DASMCs in our previous transcriptomic study, a study in which NDUFS2 was uniquely downregulated by normoxia, amongst all other Complex I subunits^28,29^. This indicates a similar effect of NDUFS2 knockdown and changing oxygen conditions to mimic fetal and neonatal environments^28^. The loss of NDUFS2 upon prolonged oxygen exposure may explain why the DA loses its ability to respond to oxygen in the days after initial closure.

## 5. Limitations

We established a key function of NDUFS2 in the DA’s oxygen response, though some questions remain. NDUFS2 has been shown to mediate the oxygen responses of other specialized oxygen responsive cells in adult humans and animals, including signalling that initiates HPV in resistance pulmonary artery smooth muscle cells^27^ and the hypoxic ventilatory response in type I cells of the carotid body^26,79^. We previously demonstrated that Complex I generation of ROS at the I_Q_ site plays a key role early in the oxygen response pathway of rabbit DASMC, with S1QEL treatment inhibiting *ex vivo* DA constriction and *in vitro* oxygen-induced increase in intracellular calcium^25^. NDUFS2 may be either a direct oxygen sensor or a participant in the sensor mechanism through its ability to modulate ROS production in proportion to PO_2_. Indeed we have extensively published that mitochondrial-derived ROS are the redox signalling mediators in mammalian oxygen sensing^17^.

In some cell lines (each line being from an individual infant), there was a complete elimination of oxygen response with knockdown of NDUFS2 (**Figure S2B**); however, across the pooled data from n=10 human DASMC cell lines studied, we did not see complete inhibition of oxygen-induced increases in intracellular calcium. This may reflect inter-individual differences in the importance of NDUFS2, as each cell line was from a different baby, with various congenital heart diseases and collected at different ages^28^. With limited patient data and a heterogeneous sample group it is difficult to conclude if NDUFS2 is the universal oxygen sensor of the DASMC or if other subunits also contribute to the DA’s oxygen response in some infants. Future studies manipulating NDUFS2 in additional human whole DA and DASMCs, as well as pharmacological manipulation in preclinical models, may further elucidate the role of NDUFS2 in DA oxygen sensing. As PDA is primarily a disease of prematurity, future studies should explore the maturational changes in NDUFS2 to examine its role in DA closure and PDA pathophysiology.

## 6. Conclusions

Identification of NDUFS2 as the oxygen sensor in the human DA advances our understanding of the molecular mechanisms underlying the physiological response of this vessel to the transition from the hypoxic fetal environment to a normoxic neonatal environment. Further exploration and manipulation of NDUFS2 and the oxygen sensing pathway may lead to improved modulation of DASMC constriction and thus DA patency. While current therapies are effective at maintaining DA patency in cases of ductus-dependent congenital heart diseases, cyclooxygenase inhibitors are ineffective in closing PDA in approximately 30% of cases, with much lower efficacy in infants born under 1000g^80^. Current pharmacologic therapy to close PDA is also associated with toxicity, including renal failure (11%), necrotizing enterocolitis (3%), and gut perforation (8%), further highlighting a need for advancement in treatments^81^. Targeting NDUFS2 in the DA oxygen sensing pathway may be a promising novel therapeutic strategy to close PDA or maintain DA patency in ductus-dependent congenital heart defects.

## Supporting information

Supplementary Transcriptomic Results

## Supplemental Material

Supplemental Methods

Table S1-S3

Figures S1-S17 Transcriptomic Results

## Data availability

For the transcriptomics, the datasets generated during and/or analyzed during the current study are available in the supplemental materials (“Supplement_TranscriptomicResults.xlsx”), with raw and processed data available through NCBI’s Gene Expression Omnibus (GEO) repository (GSE335748, https://www.ncbi.nlm.nih.gov/geo/query/acc.cgi?acc=GSE335748). Any other datasets generated during and/or analyzed during the current study are available from the corresponding author on reasonable request.

## Patient Consent

Ethics approval was obtained at each university where the samples were collected. Human DA samples were isolated during the course of congenital heart surgery at either University of Chicago (IRB number A3523–01) or University of Nebraska (IRB number 100–11-EP). Informed consent for the congenital heart surgery was obtained from the parents by the cardiovascular surgeon or his team. The focus of the consent is of course the risks and benefits of the surgery itself. However, the parents are also told that the ductus may be used in a research study to understand the mechanism by which the ductus normally closes at birth. As with all consent forms, the parents are told that their infant’s participation in the research aspects of the procedure are entirely optional. Consent is obtained in writing and is maintained by the surgeon in the medical record. The ductus is only provided to us if the cardiovascular surgeon determines that the ductus or part of the ductus is to be resected. To avoid risk to issues of confidentiality we do not review the subjects’ charts, attend the surgery, or keep a consent form.

## Author Contributions

RB, KDS, JM, and SA made substantial contributions to conception and design; RB, AM, JM, AD, KHC, BO, TN, and CCTH made substantial contributions to acquisition of data, or analysis and interpretation of data. All authors contributed to drafting the article or revising it critically for important intellectual content. All authors have provided final approval of the version to be published.

## Sources of Funding

This work was supported by a grant from the Canadian Institutes for Health Research (CIHR 183762).

## Competing Interests

None

## Notes

### Competing Interest Statement

The authors have declared no competing interest.

### Summary of Updates

New authors added; Figure 4 and 5 combined; expanded heat map in Figure 6 (now Figure 5) from 20 to 50 differentially expressed genes; Figure 7 (now Figure 6) updated with addition of panel D; new Figure 7 for comparison of transcriptomic effects of electron transport chain Complex I knockdowns; Supplemental files updated; discussion updated to clarify bioenergetic compensation with knockdown of essential electron transport chain subunits.

