## Supplementary Methods and Figures for "The Core Subunit NDUFS2 in Mitochondrial Complex I is Critical to Oxygen Sensing in Human Ductus Arteriosus Smooth Muscle Cells"

### 1. Supplementary Methods

#### 1.1. Ethics Approval

Previously isolated human DASMC were used. Ethics approval was obtained at each institution where the research was conducted. Queen's University ethics approval was obtained from Health Sciences and Affiliated Teaching Hospitals Research Ethics Board (HSREB) for the continued use of the human cell lines in this research (TRAQ # 6007784).

#### 1.2. Human DASMC Cell Culture

All cell culture reagents were acquired from Thermo Fisher Scientific (Mississauga, ON, Canada). Primary cell lines were previously established from surgically harvested DAs of infants undergoing congenital heart surgery as previously described[1]. Cell lines were previously purified using fluorescence activated cell sorting and phenotypically confirmed using flow cytometry[2]. hDASMC were grown in Human Vascular Smooth Muscle Cell Basal Medium, supplemented with 5% Smooth Muscle Growth Supplement, 10% Fetal Bovine Serum, 1% L-Glutamine, 1% penicillin/streptomycin, and 10μg/mL ciprofloxacin HCl. Cells were used within the first five passages and were carefully maintained in hypoxia (2.5% $O_2$, 5% $CO_2$, balance $N_2$) prior to experimental normoxic exposure (19.6% $O_2$, 5% $CO_2$, balance $N_2$).

#### 1.3. Silencing RNA (siRNA) Knockdown

Silencing RNAs targeted at NDUFS1, NDUFS7, UQCRFS1, COX4I2, and Negative Control (Cat# 51-01-14-04) were obtained from Integrated DNA Technologies (IDT, Coralville, IA) and silencing RNA targeted at NDUFS2 was obtained from Qiagen (Hilden, Germany). The sequences for the siRNA used are shown in **Table S1**.

Cells were seeded one day prior to siRNA treatment into 35mm dishes or 6-well plates. siRNAs were transfected with Lipofectamine™ RNAiMAX Transfection Reagent (Cat#13778150, ThermoFisher Scientific, Mississauga, ON, Canada**)** following the manufacturer's protocol.

Four siRNAs targeting NDUFS2 and three siRNAs targeting NDUFS7 were screened by measuring the knockdown efficiency using qPCR following 48-hours treatment. The siRNA for each knockdown condition was selected as any that depleted at least 80% of mRNA and/or the one that produced maximal mRNA knockdown. The optimal time-point of siRNA knockdown was tested in two cell lines at 24-, 48- and 72-hours via Western Blot.

**1.4.mRNA Quantification with Quantitative Real Time PCR (qPCR)**

Cells treated with siRNA were collected and lysed in TRI Reagent ® (Cat# T9424-100ML, Lot # MKCK9023, Sigma-Aldrich, Oakville, ON, Canada) and RNA was extracted using the Direct-zol™ RNA Miniprep kit (Cat# R2050, Zymo Research, California) following the manufacturer's protocol. RNA was quantified using the Qubit™ RNA High Sensitivity (HS) Assay (Cat# Q32852, ThermoFisher Scientific). First-strand cDNA synthesis was completed using qScript cDNA SuperMix (QuantaBio, Beverly, MA), following the manufacturer's protocol, with 300µg RNA input per sample.

qPCR was conducted using TaqMan™ probes (ThermoFisher Scientific) and PerfeCTa Fastmix II (no ROX, Quantabio) according to manufacturer's protocol, using the QuantStudio™ 3 – 96-well 0.2mL Block (cat#A28567, ThermoFisher Scientific). The internal control in all experiments was the housekeeping gene Eukaryotic 18S rRNA Endogenous Control (Cat#4319413E, ThermoFisher Scientific). The following primers were used to compare expression in cells treated with specific versus control siRNA: *NDUFS2* (Hs01035077_g1),

*NDUFS1* (Hs00192297_m1), *NDUFS7* (Hs01086223_m1), *UQCRFS1* (Hs0419251_g1), and *COX4I2* (Hs00261747_m1). Samples were run in triplicate, averaging the $C_T$ values, using a no template control well (water instead of RNA input) as the negative control. The delta $C_T$ ($\Delta C_T$), difference between the target gene and the housekeeping gene (18S rRNA), was calculated for all targets and differences between $\Delta C_T$ in siControl treatment and gene targeting siRNA were compared using two-tailed unpaired t-tests. Gene expression data from qPCR validation is all reported as log2 fold change[3].

**1.5.Protein Quantification of ETC Knockdown with Immunoblotting**

Protein was extracted with cell lysis buffer (Cat# 9803S, Cell Signaling Technologies, Beverly, MA) and quantified using the Pierce™ BCA protein assay (Cat# 23225, ThermoFisher Scientific) SpectraMax M3 (Molecular Devices, San Jose, CA). Cell lysates (20-50µg) were denatured with 2-mercaptoethanol (Cat#M3148, Sigma-Aldrich) and NuPAGE™ LDS Sample Buffer (Cat#NP0007, ThermoFisher Scientific). Samples were analyzed for immunoblot analyses on 4-12% NuPAGE gels (Life Technologies, Carlsbad, CA). The proteins were detected using the indicated antibodies and ECL-Plus Western Blotting Detection System (GE Healthcare, Piscataway, NJ), as previously described[4]. For detection of ETC subunits, the following antibodies were used: NDUFS1 (Ab169540, Abcam, Toronto, ON, Canada), NDUFS2 (Ab192022, Abcam), NDUFS7 (Cat# 15728-1-AP, ThermoFisher Scientific), UQCRFS1 (NBP1-87826, Novus Biologicals, Centennial, CO), and COX4I2 (Ab33985, Abcam). All membranes were stripped with Restore™ Western Blot Stripping Buffer (Cat# 21059, ThermoFisher Scientific) and probed for beta-actin as a loading control (Cat#A5441, Sigma-Aldrich). Densitometry was quantified using Fiji software[5], with each target normalized to beta-actin and compared to untreated and siControl treated cells.

##### 1.6. Confocal Live Cell Imaging

Three days prior to imaging, hDASMC were plated into 35mm glass bottom dishes (No. 1.5 uncoated γ-irradiated, P35G-1.5-14-C MatTek Corporation, Ashland, MA) in Human Vascular Smooth Muscle Cell supplemented media at a density of 1 x $10^5$ cells per dish. The cells were treated with siRNA the day after plating. Calcium imaging experiments were conducted 48-hours after siRNA treatment in an oxygen-sensing assay buffer with the dye Cal-520 (ab171868, Abcam), as previously described[2]. All live cell imaging was performed using a a Leica TCS SP8 X confocal microscope (Leica Microsystems, Wetzlar, Germany). Mitochondrial morphology experiments were conducted 48-hours after plating in an oxygen sensing assay buffer with the dye TMRM (tetramethylrhodamine methyl ester, T668, ThermoFisher Scientific). Dishes were stained with 2µM TMRM in growth media for 30 minutes at 37°C, washed with assay buffer and imaged in 1mL assay buffer for 10 minutes in hypoxia (3% $O_2$, 5% $CO_2$, balance $N_2$) followed by 10 mintues normoxia (room air), with mitochondrial network and fragmentation quantified with machine learning as previously described[6]. Mitochondrial ROS imaging experiments were also conducted 48-hours after siRNA treatment in an oxygen-sensing assay buffer with the dye MitoROS 580 (ab219943, Abcam), and using 1-hour pretreatment with 50µM mitochondrially-targeted antioxidant (MitoTEMPO, SML0737, Sigma-Aldrich) as a positive control, as previously described[7]. Cells were imaged during hypoxia (3% $O_2$, 5% $CO_2$, balance $N_2$) or normoxia (room air), with care taken to avoid oxygen exposure except when administering a normoxic challenge. Cells were imaged for 20-minutes in hypoxia, followed by 20-minutes in normoxia, capturing one frame every 15-seconds in a tile-scan of four fields of view. During calcium imaging experiments, the cells were challenged with 80mM potassium chloride (KCl) while imaging to act as a positive control by depolarizing the smooth muscle cell membrane. Cells and dishes that did not have a

KCl response ($\geq$2-fold increase in intensity above hypoxic baseline) were excluded from analysis. The oxygen-induced calcium response was measured as the increase from hypoxic baseline. Cell constriction was also measured to determine oxygen-responsiveness. The length of ten randomly selected cells were measured during the hypoxic baseline and during the 20-minutes of normoxia. The cell constriction is reported as percentage cell shortening relative to the hypoxic baseline.

##### 1.7.Seahorse Micropolarimetry

Mitochondrial metabolism was quantified by measuring the oxygen consumption rate (OCR) using an XFe24 extracellular flux analyzer (Agilent, Santa Clara, CA), as previously described[7,8]. Assays were completed 48 hours or 96 hours after siRNA treatment, with cells plated at 40,000 cells per well the day prior to the experiment in a 24-well Xfe24 cell culture microplate (Agilent), following the manufacturer's protocol for adherent cells. Seahorse XF base medium (Cat#103335-100, Agilent) was supplemented with 10mM D-Glucose (Cat# G8270, Sigma-Aldrich), 1mM Sodium Pyruvate (Cat#11360-070, ThermoFisher Scientific), and 2mM L-glutamine (Cat# SH3003401, ThermoFisher Scientific). The Seahorse XF Cell Mito Stress Test was used to assess metabolic function, adding first oligomycin (Cat# O4876, Sigma-Aldrich), then carbonyl cyanide-4 (trifluoromethoxy) phenylhydrazone (FCCP, C2920, Sigma-Aldrich), and finally rotenone in combination with Antimycin A (Cat# R8875 & Cat# A8674, Sigma-Aldrich) while measuring OCR. All OCR values were normalized to total protein content in each well following protein quantification using the Pierce™ BCA protein assay.

##### 1.8.Complex I Activity Assay

For the determination of Complex I activity following ETC subunit knockdown via 48-hours siRNA treatment, a Complex I activity dipstick assay (ab109720, Abcam) was used, following the manufacturer's protocol. 25µg protein was loaded for each sample per reaction, and dipsticks were developed for 45-minutes. Densitometry of the immunocaptured Complex I band was quantified using Fiji[5], with knockdown conditions compared to siControl treated cells.

##### 1.9.Complex III Activity Assay

For the determination of Complex III activity, mitochondria were isolated from hDASMC treated for 48-hours with siRNA using the Mitochondria Isolation Kit for Cultured Cells (ab110170, Abcam) following manufacturer's protocol. 1.5µg isolated mitochondria per siRNA-treated hDASMC condition used for to assess Complex III activity with the mitochondrial Complex III activity assay kit (ab287844, Abcam), following the manufacturer's protocol. Absorbance at 550nm of samples with and without Complex III inhibitor antimycin A was measured at 30-second intervals for 10-minutes following addition of oxidized cytochrome c. The concentration of reduced cytochrome c of samples over time was calculated based on the standard curve, with the net Complex III activity determined as the difference in activity with and without Antimycin A.

##### 1.10. Complex IV Activity Assay

Complex IV activity was assessed using the Complex IV human enzyme activity microplate assay kit (ab109909, Abcam), following the manufacturer's protocol. 25µg of each sample was loaded into the microplate, and sample only (no assay buffer) served as the background control. The absorbance of the plate was measured at 550nm for 80-minutes every minute. Complex IV activity was calculated as the rate of oxidation of cytochrome c, seen as a decrease in absorbance (slope), subtracting the absorbance in background control wells.

##### 1.11. RNA Sequencing of siRNA-Treated Cells

3' RNA sequencing was conducted on five hDASMC cell lines grown in hypoxia, each treated for 48 hours with siRNA targeting negative control, NDUFS1, NDUFS2, NDUFS7,

UQCRFS1, or COX4I2. RNA was extracted and samples prepared for sequencing using the QuantSeq 3' FWD mRNA-Seq Library Prep Kit for Illumina (Lexogen, Austria), as previously described[2], with 130 ng RNA as the input.

The RNA sequencing analysis was conducted using the DESeq2 (V 1.44.0) package in R (V 4.4.0)[9,10]. Each knockdown condition was compared to the siControl condition, with DEGs identified at an alpha of 0.05 significance level after correcting for multiple testing with the Benjamini & Hochberg method[11]. The top 50 DEGs, chosen by absolute value of log fold change, between siNDUFS2 and siControl (14 upregulated and 6 downregulated with siNDUFS2) were compared across knockdown conditions, creating heatmaps with the pheatmap package (V 1.0.13). Utilizing the gprofiler2 package (V 0.2.3), we performed enrichment analysis through Gene Ontology: Biological Process, Cellular Components, and Molecular Function, as well as KEGG, TF, MIRNA, WP, and REAC pathways[12,13]. The search string "mitoc" was used to examine Gene Ontology pathways relating to mitochondrial functions. Plots of top 20 Gene Ontology pathways were created using ChiPlot[14]. For the creation of heatmaps of genes within selected GO terms, DOSE (v4.6.0)[15], enrichplot (v1.32.0)[16], and clusterProfiler (v4.20.0)[16] were used. To determine the genes contained in the mitochondrial and metabolic GO pathways of the top 20 pathways uniquely regulated by siNDUFS2 (**Figure 6D**) matching the original enrichment results, an archived version of Ensembl was used to compare the genes annotated within selected GO terms from "Ensembl 111, Ensembl Genomes 58 (database built on 2024-01023)" (https://biit.cs.ut.ee/gprofiler/page/archives) to the 578 genes uniquely regulated by siNDUFS2. For comparisons between siNDUFS2 and siNDUFS7, and siNDUFS2 and siNDUFS1 we performed enrichment analysis for Gene Ontologies using the gprofiler2 package (V 0.2.4). The top 20 pathways for comparisons between siNDUFS2 and siNDUFS7, and siNDUFS2 and siNDUFS1 were the 20 most significantly enriched pathways (smallest p-values), with the lists then ordered by number of genes in each GO term (**Figures S13 and S16**).

##### 1.12. Statistics

All data analysis was conducted using Prism 10 (GraphPad Software, LLC, San Diego, CA). Data are presented as mean ± standard error mean (SEM) unless otherwise stated. Descriptive statistics were performed to check for equal variance between groups and the Shapiro-Wilk test was used to test for normal distribution. The ROUT method was used to test for and remove outliers[17]. Paired t-tests were used to compare the knockdown efficiency between siControl treated cells and target siRNA, for both qPCR and immunoblot quantification. For individual cell line data, the Friedman test with Dunn's multiple comparisons test was used to test the calcium response under each knockdown condition, comparing Cal-520 signal in hypoxia, normoxia, and with KCl treatment. When comparing normoxic response across knockdown conditions across all cell lines, Kruskal-Wallis with Dunn's multiple comparisons test was used. The same test was used to compare cell length in hypoxia versus normoxia to quantify DASMC constriction. A linear mixed effect model, using the Satterthwaite approximation, of untreated, siControl, siNDUFS2, and MitoTEMPO treated dishes was used to quantify differences in mitochondrial network between hypoxia and normoxia. The Friedman test with Dunn's multiple comparisons test was used to compare the parameters from the Seahorse Mito Stress Test across knockdown conditions (basal respiration, proton leak, maximal respiration, spare respiratory capacity, spare respiratory capacity as percent of basal, non-mitochondrial respiration, ATP-linked respiration, and coupling efficiency). A repeated-measured one-way ANOVA with Dunnett's multiple comparisons test, with a single pooled variance was used to compare Complex I activity in all knockdown conditions compared to siControl. For Complex III activity, the Friedman test with Dunn's multiple comparisons test was used, comparing all knockdown conditions to the untreated Complex III activity. For Complex IV activity, the Kruskal-Wallis test with Dunn's correction for multiple comparisons was used to compare all knockdown conditions to siControl Complex IV activity.

| | |
|---|---|
| NDUFS1 | 5’-GAAGAGUGGAUCUCUGAUAAAACCA-3’ |
| | 3’-UACUUCUCACCUAGAGACUAUUUUGGU-5’ |
| NDUFS2 | 5’-CUGGCGAAAUCGGACAAUUGA-3’ |
| | 3’-UAGACCGCUUUAGCCUGUUAACU-5’ |
| NDUFS7 | 5’-CCCGUGAGGUUGUCAAUAAACCUGC-3’ |
| | 3’-CAGGGCACUCCAACAGUUAUUUGGACG-5’ |
| UQCRFS1 | 5’-AGCAUGAUCUAGAUCGAGUAAAGAA-3’ |
| | 3’UGUCGUACUAGAUCUAGCUCAUUUCUU-5’ |
| COX4I2 | 5’CACCUUGACGGACGAGCGGAAAGC-3’ |
| | 3’GUGGAACUGCCUGCUCGCCUUUCG-5’ |

**Table S1: siRNA Sequences**

The sequences of the silencing RNAs (siRNAs) with optimal knockdown efficiency, used to selectively knock down mitochondrial electron transport chain subunits for *in vitro* studies.

| Cell Line | Sex | Age | Diagnosis |
|---|---|---|---|
| 1 | Male | 19 days | Aortic coarctation, Hypoplastic left ventricle |
| 2 | Female | 8 months | VSD, PDA, CHF/PH |
| 3 | Male | 11 days | Aortic coarctation, bicuspid aortic valve, small PDA |
| 4 | Male | 10 days | - |
| 5 | Male | - | HLHS |
| 6 | Male* | - | - |
| 7 | Male | 5 days | HLHS |
| 8 | Female | 6 days | D-TGA, PDA, PFO |
| 9 | Male | 4 days | TGA |
| 10 | Male* | - | - |
| 11 | Female | 18 days | - |

**Table S2: Human DA Donor Demographics**

All available demographics of the infants from whom the DASMC were isolated are listed. Missing data (age and/or diagnosis) are indicated by dashes (-). VSD: ventricular septal defect, PDA: patent ductus arteriosus, CHF: congestive heart failure, PH: pulmonary hypertension, HLHS: hypoplastic left heart syndrome, (D-)TGA: (dextro-) transposition of great arteries, PFO: patent foramen ovale. * indicates sex determined by qPCR of X-inactivation gene XIST[22]. Adapted with permission from Bentley et al. 2021, Genomics 113[22].

| Figure | Cell Lines Used |
|---|---|
| **Figure 2** | **A)** 1, 5, 6, 10 |
| | **B)** 1, 3, 5, 6, 9, 10 |
| | **C)** 1, 5, 6, 8, 9, 10 |
| | **D)** 5, 6, 8, 9, 10 |
| | **E)** 1, 5, 6, 8, 9, 10 |
| | **F)** 1, 3, 5, 6, 10 |
| | **G)** 1, 5, 6, 8, 9, 10 |
| | **H)** 1, 5, 6, 8, 9, 10 |
| | **I)** 5, 6, 10 |
| | **J)** 1, 5, 6, 9 |
| **Figure 3** | **B)** 4, 5, 8, 9, 10 |
| | **D/E)** 1, 2, 3, 4, 5, 7, 8, 9, 10, 11* |
| | **G/H)** 5, 8, 9, 10 |
| **Figure 4** | **B)** 1, 2, 3, 5, 6, 8, 9, 10 |
| | **C)** 2, 3, 4, 6, 10 |
| | **D)** 1, 2, 3, 5, 6, 8, 9, 10 |
| | **F/G)** 1, 3, 5, 8, 9, 10 |
| | **H/I)** 4, 5, 9, 10 |
| **Figure 5-7** | 4, 5, 6, 7, 11 |

**Table S3: Index of Cell Lines Used per Experiment**

List of cell lines used in each assay result, indicated by Figure and panel letter. Cell line number corresponds to donor demographics in Table S3. *Cell line conditions excluded due to lack of KCl response: Untreated = 5, 10; siNDUFS1 = 5; siNDUFS2 = 7, 8; siNDUFS7 = 1, 2; siUQCRFS1 = 4, 7, 8; siCOX4I2 = 5, 7.

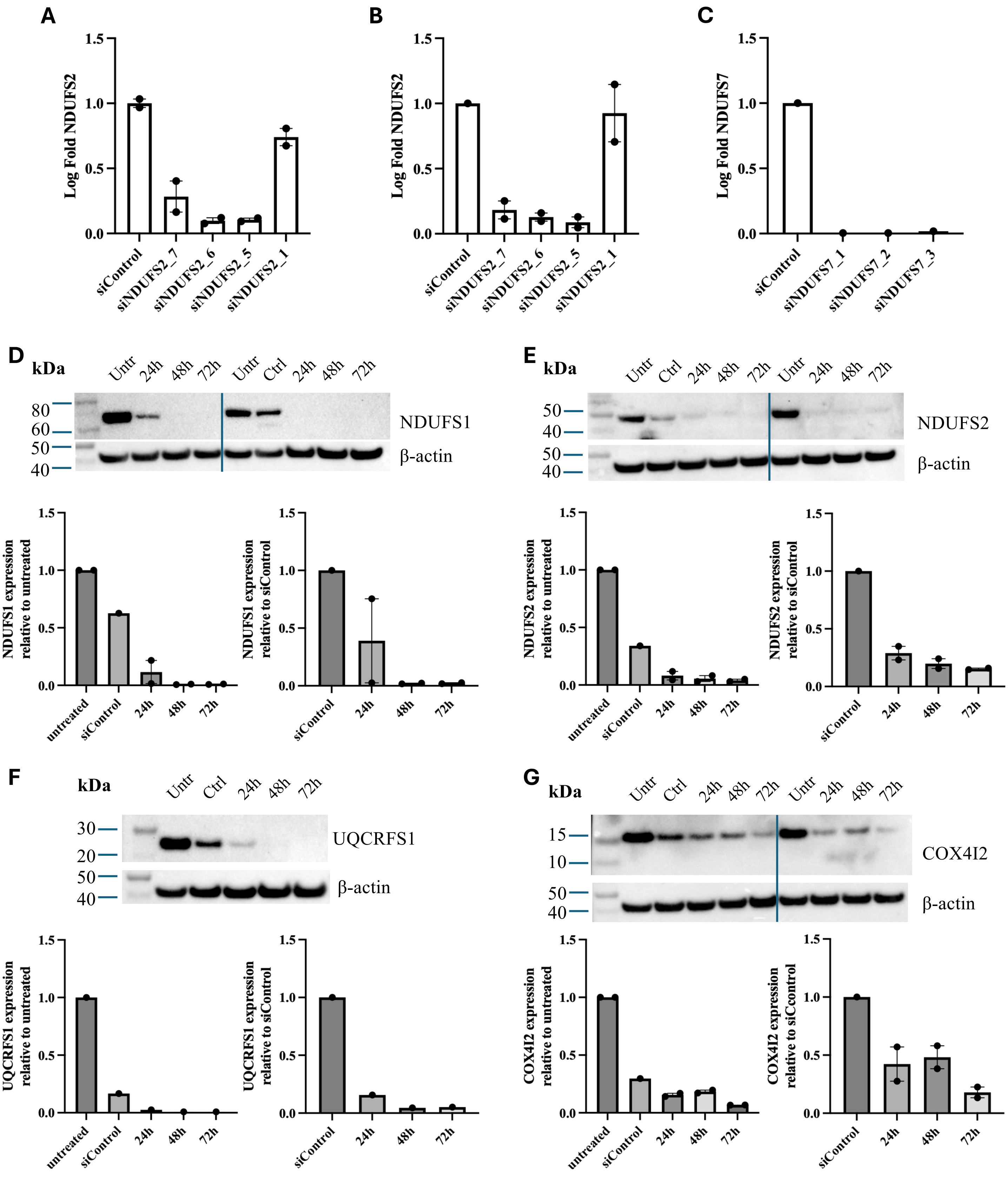


**Figure S1: Silencing RNA (siRNA) Determined Optimal Knockdown of Electron Transport Chain (ETC) Subunits**

Four siRNAs targeting NDUFS2 were screened following 48-hours treatment in hypoxia (**A**) or normoxia (**B**) (n=2 cell lines); greatest knockdown was achieved with siNDUFS2_5 versus siControl, with siNDUFS2_5 used for all further studies. **C**) Three siRNA targeting NDUFS7 were screened following 48-hours treatment (n=1 cell line), with siNDUFS7_1 chosen for further experiments. Protein knockdown efficiency at 24-, 48-, and 72-hours was assessed via western blot for siRNA targeting NDUFS1 (**D**), NDUFS2 (**E**), UQCRFS1 (**F**), and COX4I2 (**G**). Protein expression was compared to untreated (no siRNA) and siControl-treated cells, with all expression normalized to β-actin. Maximal knockdown occurred at 48-hours with siNDUFS1, siNDUFS2, and siUQCRFS1; maximal knockdown of COX4I2 occurred at 72-hours.

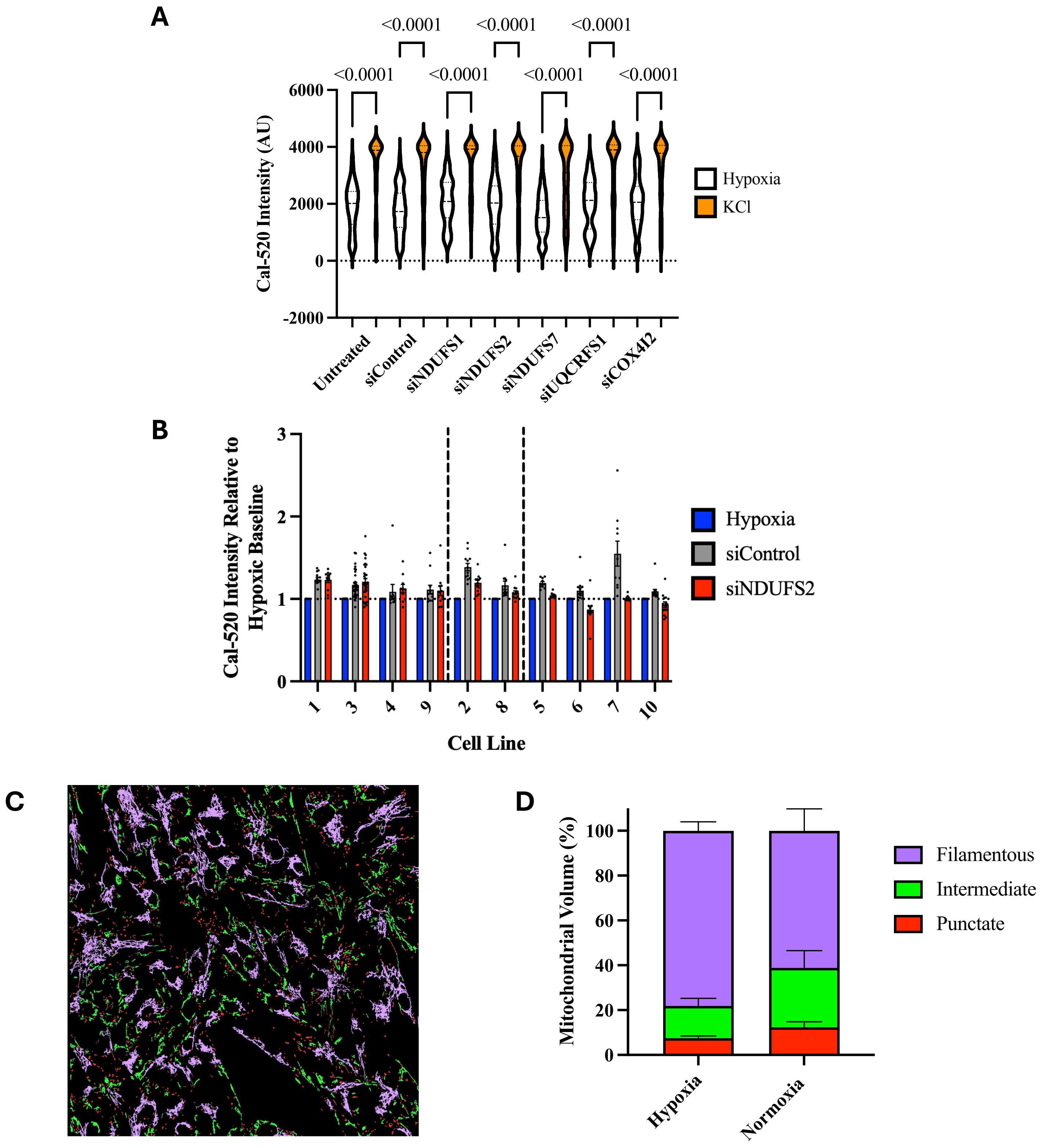


**Figure S2: hDASMC Oxygen Responsiveness Quantification via Confocal Imaging**

**A**) Cal-520AM was used to measure changes in intracellular calcium with live confocal imaging (n=10 cell lines). Potassium chloride (KCl) was used as a positive control for DASMC depolarization and rise in intracellular calcium, with cells lacking a KCl-induced rise in intracellular calcium excluded from analysis. siRNA treatments did not decrease DASMC KCl response. **B**) Oxygen responsiveness, quantified as Cal-520 intensity in normoxia relative to hypoxia (normoxia/hypoxia), with 48-hours siControl or siNDUFS2 reveals heterogeneity in effect strength among cell lines. There were four cell lines where there was no effect of siNDUFS2 versus siControl on oxygen response, two cell lines with moderate depression in oxygen responsiveness with siNDUFS2, and four cell lines where siNDUFS2 completely eliminated oxygen response, with no change or reduction in intracellular calcium from hypoxia to normoxia. **C**) Live cell imaging with mitochondrial dye TMRM (tetramethylrhodamine methyl ester) was used to characterize and quantify the mitochondrial network of hDASMC under hypoxia and normoxia, with machine learning classifying mitochondria as punctate, intermediate or filamentous. **D**) The percentage of punctate, intermediate, and filamentous mitochondria per field of view was quantified under hypoxia and following 10 minutes normoxia (n=5 cell lines).

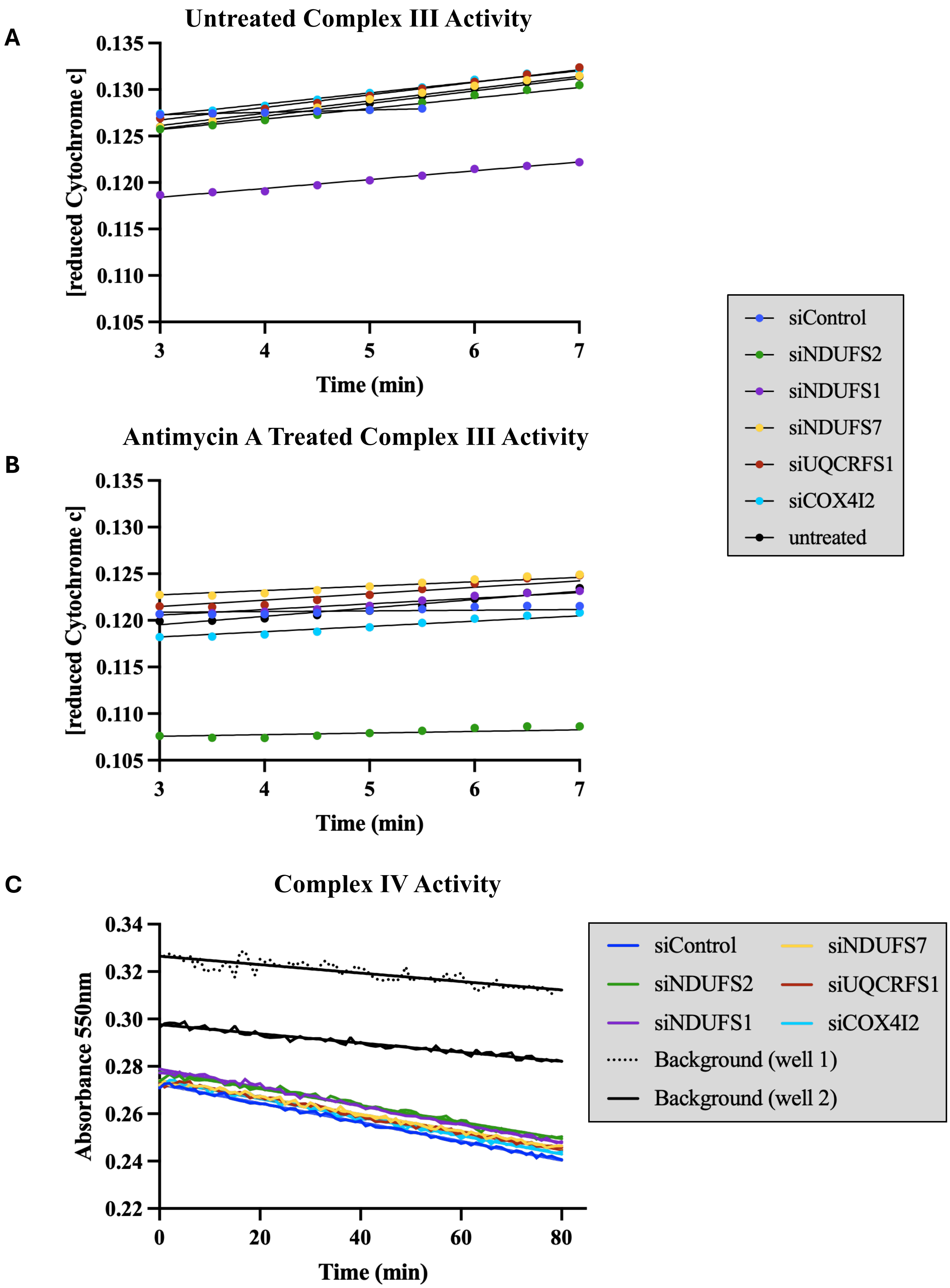


**Figure S3: Raw Complex III and IV Activity Assay Data**

Raw data from one human DASMC cell line, from which Complex III and IV activity were derived. **A**) Complex III activity was measured as change in reduced cytochrome c over time untreated and (**B**) with antimycin A treatment. **C**) Complex IV activity was measured as change in absorbance at 550nm over time, adjusted to background.

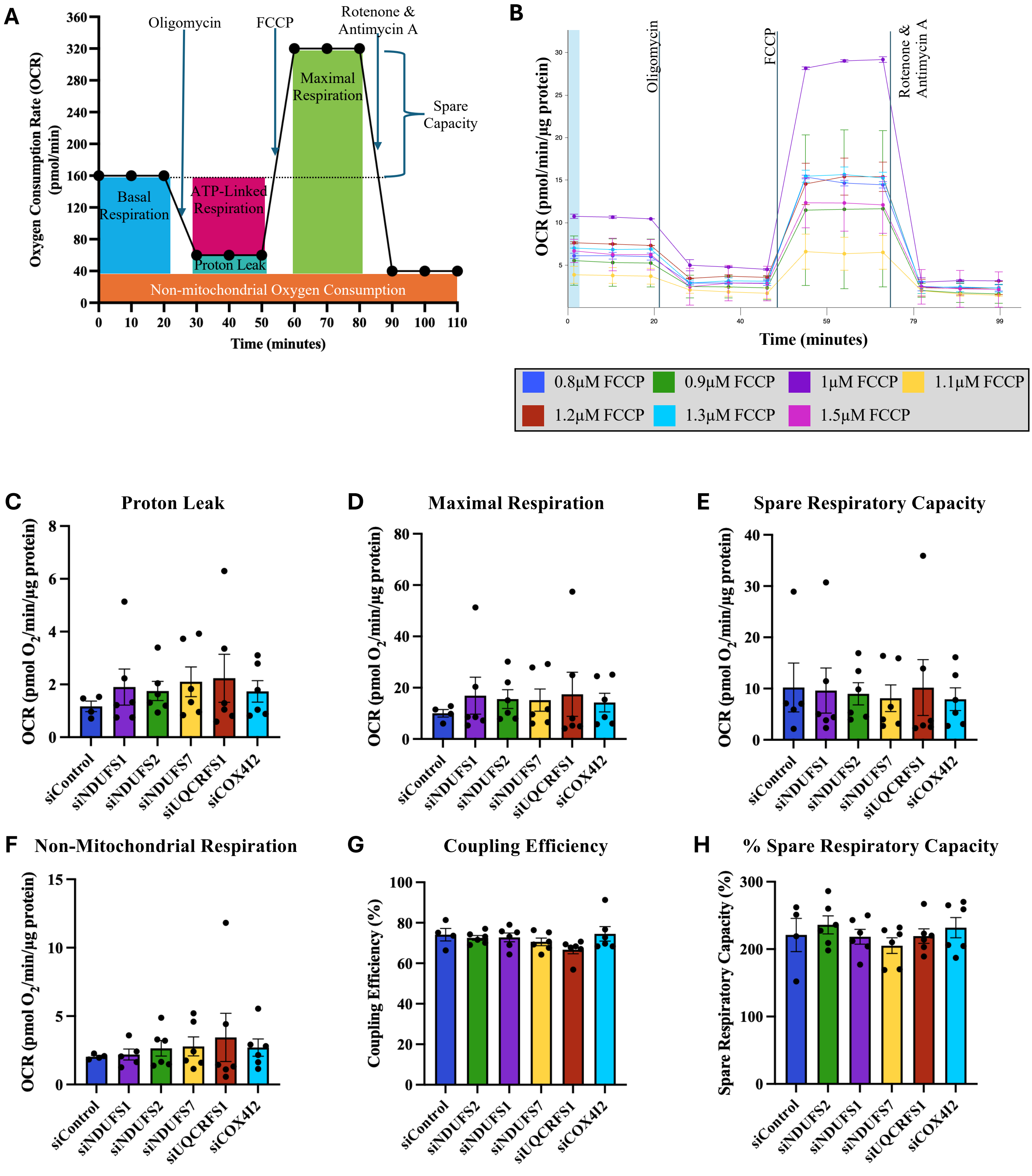


**Figure S4: Mito Stress Test Optimization and Additional Mitochondrial Metabolic Parameters**

High-precision respirometry "Mito Stress Tests" were run on an Agilent XFe24 Analyzer. **A**) The schematic demonstrates the sequence of drug additions and derivation of metabolic parameters. **B**) Experimental optimization used 1.2µM oligomycin, and 1µM each rotenone and antimycin A, based on prior published work. The optimal concentration of FCCP was experimentally determined, testing seven FCCP concentrations ranging from 0.8-1.5µM. Highest maximal respiration was observed with 1µM FCCP. Knockdown of ETC subunits had no effect on **C**) proton leak, **D**) maximal respiration, **E**) spare respiratory capacity, **F**) non-mitochondrial respiration, **G**) coupling efficiency, or **H**) spare respiratory capacity as a percent of basal respiration. OCR was normalized to total µg protein. FCCP = carbonyl cyanide-p-trifluoromethoxyphenylhydrazone.

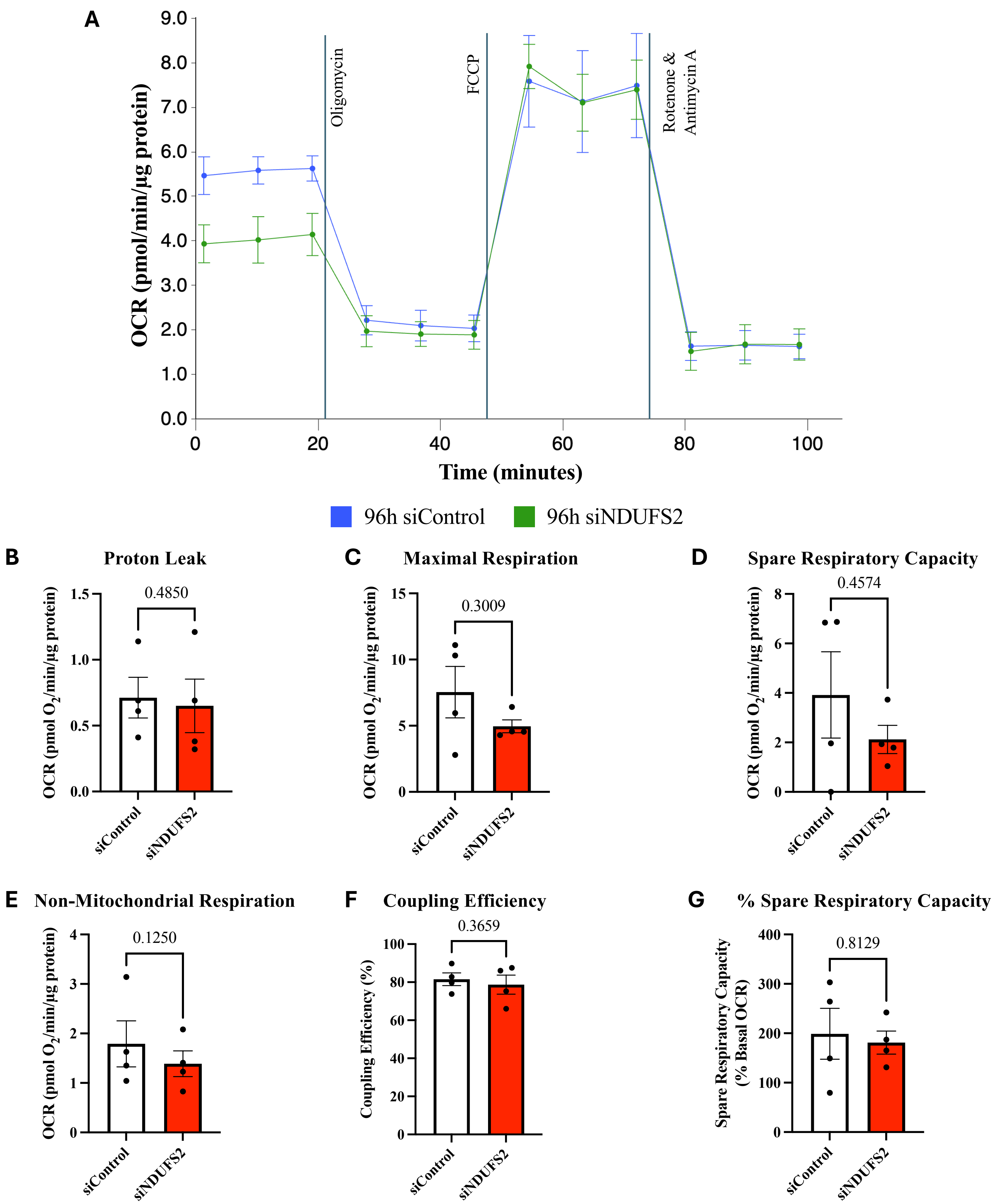


**Figure S5: Mito Stress Test Demonstrates No Significant Effect of 96-hours NDUFS2 Knockdown on Additional Mitochondrial Metabolic Parameters**

High-precision respirometry “Mito Stress Tests” were run on an Agilent XFe24 Analyzer. **A**) Representative trace of Mito Stress Test following 96-hours treatment of hDASMC with siControl or siNDUFS2, run with 1.2µM oligomycin, 1µM FCCP, and 1µM each rotenone and antimycin A. Knockdown of NDUFS2 had no significant effect on **B**) proton leak, **C**) maximal respiration, **D**) spare respiratory capacity, **E**) non-mitochondrial respiration, **F**) coupling efficiency, or **G**) spare respiratory capacity as a percent of basal respiration (n=4 cell lines, 5 wells per condition). OCR was normalized to total µg protein. FCCP = carbonyl cyanide-p-trifluoromethoxyphenylhydrazone.

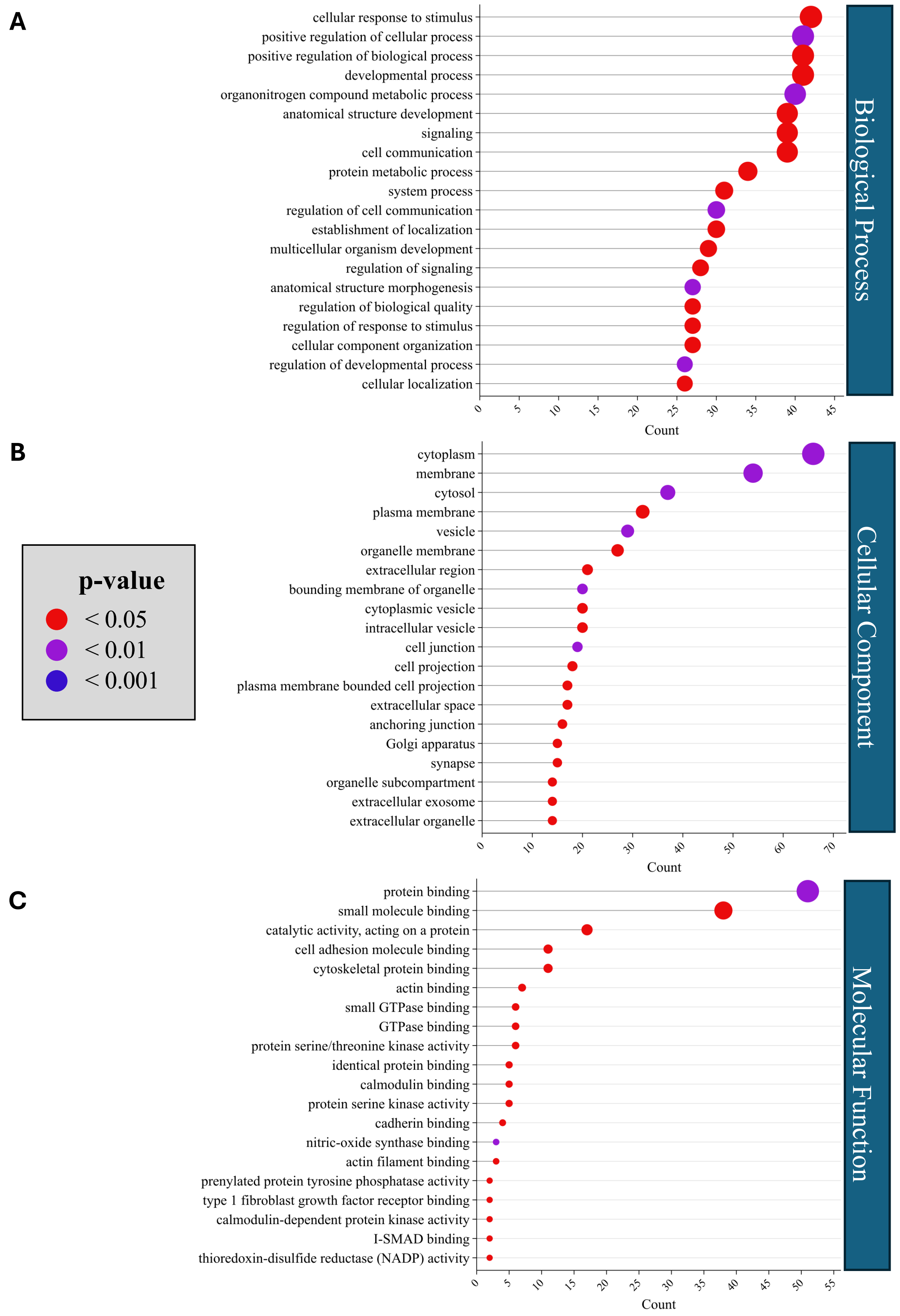


**Figure S6: Gene Ontology (GO) Enrichment Analysis of Commonly DEGs Across ETC Knockdowns**

The top 20 significantly enriched (adjusted p<0.05) GO terms of 677 commonly DEGs between siControl and each ETC knockdown condition (siNDUFS1, siNDUFS2, siNDUFS7, siUQCRFS1, and siCOX4I2) in the **A**) Biological Process, **B**) Cellular Component, and **C**) Molecular Function domains. The common DEGs primarily relate to developmental processes (GO:0032502, GO:0048856, GO:0007275, GO:0009653, GO:0050793), signaling (GO:0051716, GO:0023052, GO:0007154, GO:0010646, GO:0023051, GO:0048583), cell adhesion (GO:0030054, GO:0042995, GO:0120025, GO:0070161), and cytoskeletal binding (GO:0008092, GO:0003779, GO:0005516) GO pathways. They also related to cellular localization, including cytoplasm (GO:0005737, GO:0005829), extracellular (GO:0005576, GO:0005615, GO:0070062, GO:0043230), membranes (GO:0016020, GO:0005886, GO:0031090, GO:0098588), vesicle (GO:0031982, GO:0031410, GO:0097708), and organelle (GO:0031090, GO:0005794, GO:0031984) gene ontologies, as well as catalytic pathways, such as GTPase binding (GO:0051020) and protein serine/threonine kinase activity (GO:0004674).

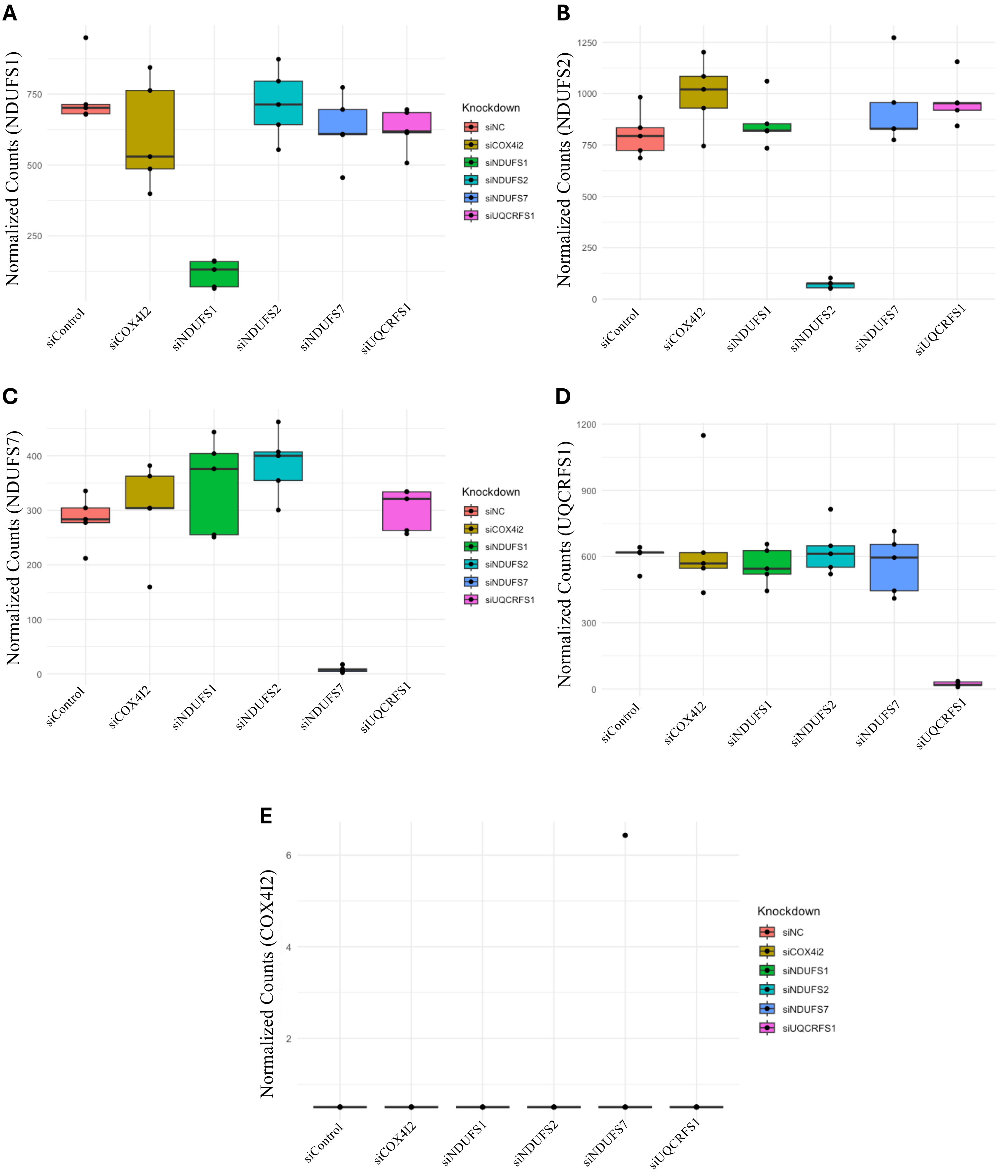


**Figure S7: Each Knockdown Did Not Affect Expression of the Other ETC Targets**

Transcriptomic results demonstrate that **A**) siNDUFS1 was the only siRNA treatment to decrease expression of NDUFS1, as shown by the decrease in normalized counts detected by 3' RNA sequencing. Similarly, **B**) siNDUFS2 was the only siRNA to decrease NDUFS2 expression, **C**) siNDUFS7 was the only siRNA to decrease NDUFS7 expression, and **D**) siUQCRFS1 was the only siRNA to decrease UQCRFS1 expression. **E**) COX4I2 was not detected in any knockdown condition.

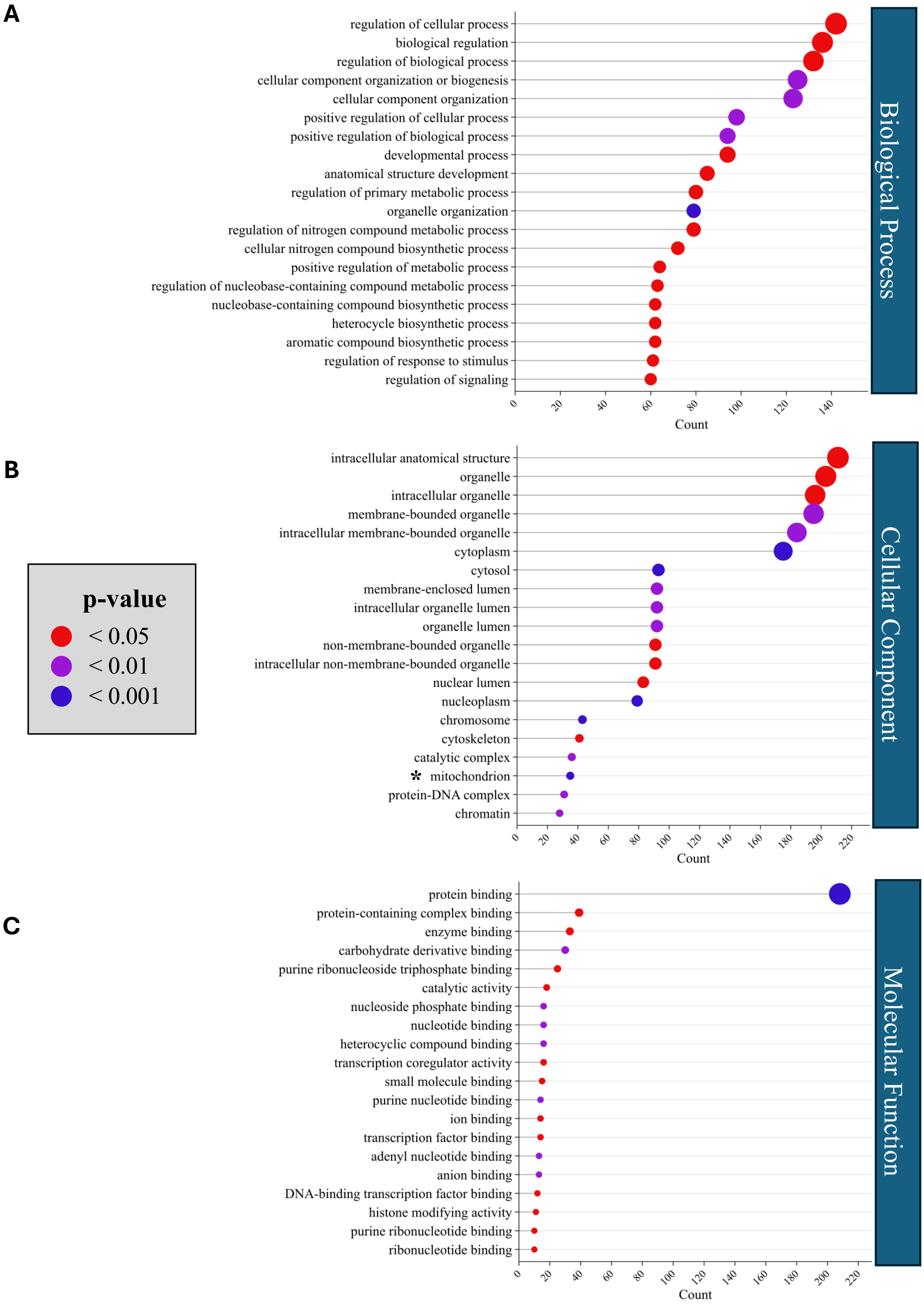


**Figure S8: Gene Ontology (GO) Enrichment Analysis of DEGs Uniquely Regulated by NDUFS1 Knockdown**

The top 20 significantly enriched (adjusted p<0.05) GO terms of the 548 genes uniquely differentially expressed between siControl and siNDUFS1 in the **A**) Biological Process, **B**) Cellular Component, and **C**) Molecular Function domains. These DEGs are primarily related to general cellular and biological process pathways, as well as nitrogen compound and nucleobase biosynthetic and metabolic processes, protein and enzyme binding, and nuclear- and organelle-related GO pathways.

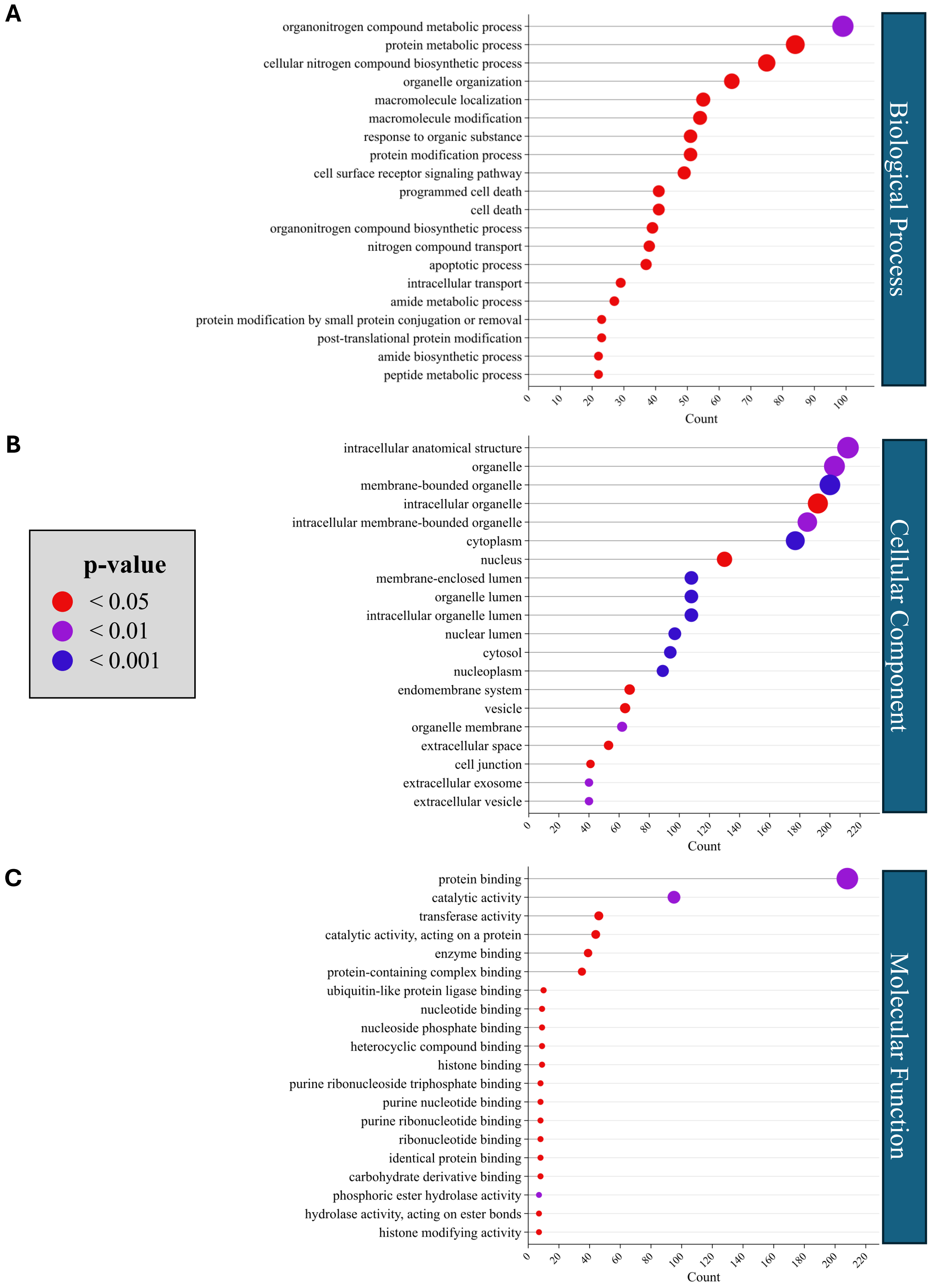


**Figure S9: Gene Ontology (GO) Enrichment Analysis of DEGs Uniquely Regulated by NDUFS7 Knockdown**

The top 20 significantly enriched (adjusted $p<0.05$) GO terms of the 765 genes uniquely differentially expressed between siControl and siNDUFS7 in the **A**) Biological Process, **B**) Cellular Component, and **C**) Molecular Function domains. The DEGs uniquely regulated by siNDUFS7 were mainly related to nitrogen and organonitrogen compound biosynthetic and metabolic process, protein and amide biosynthesis and binding, nucleotide and nucleoside binding, and various cellular components (organelle, cytoplasm, vesicle, nucleus).

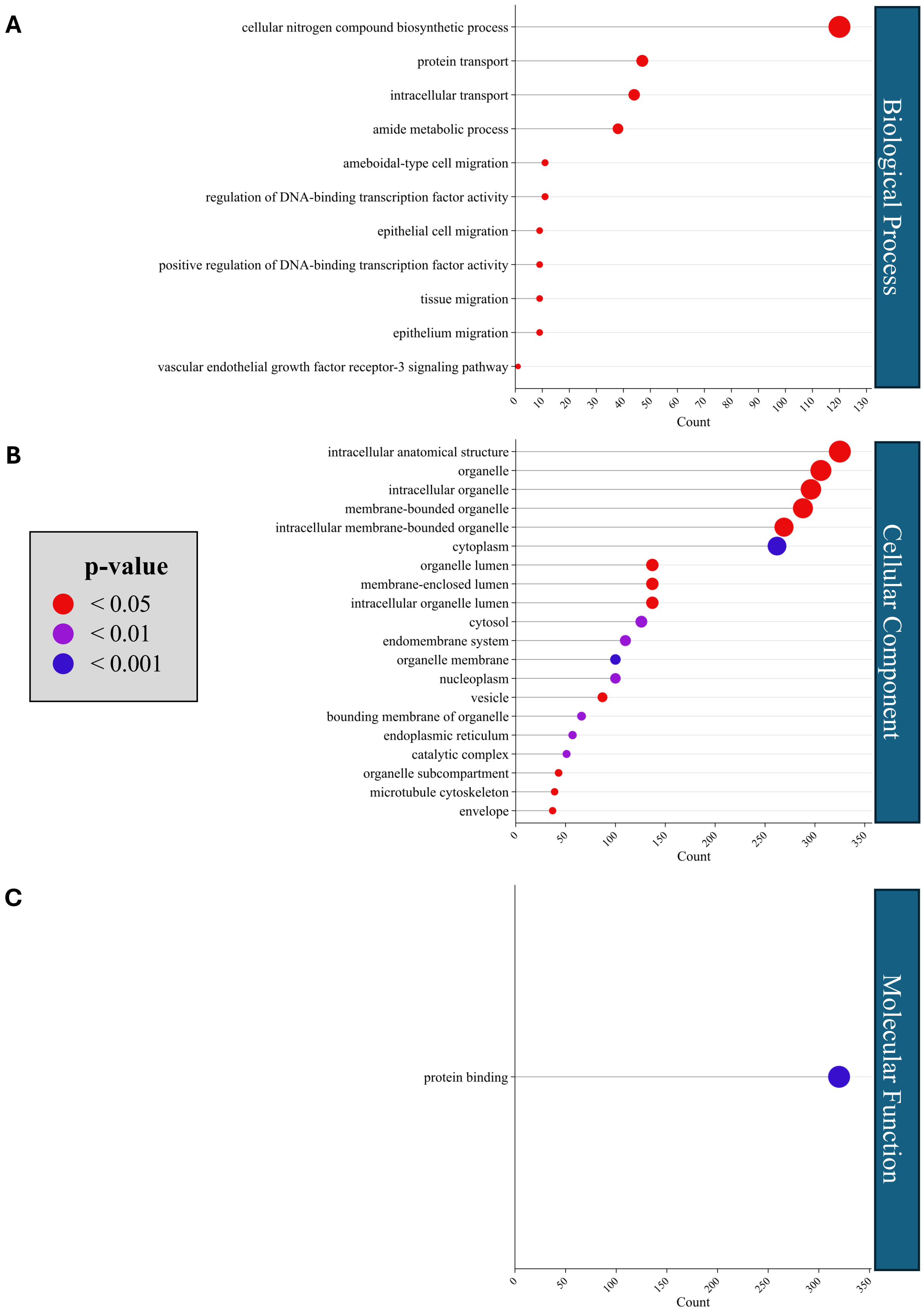


**Figure S10: Gene Ontology (GO) Enrichment Analysis of DEGs Uniquely Regulated by UQCRFS1 Knockdown**

The top 20 significantly enriched (adjusted $p<0.05$) GO terms of the 840 genes uniquely differentially expressed between siControl and siUQCRFS1 in the **A**) Biological Process, **B**) Cellular Component, and **C**) Molecular Function domains. These unique DEGs had enriched GO pathways relating to protein binding and transport, nitrogen compound biosynthetic process, cell migration, DNA-binding transcription factor activity, organelles, cytoplasm, and vesicles.

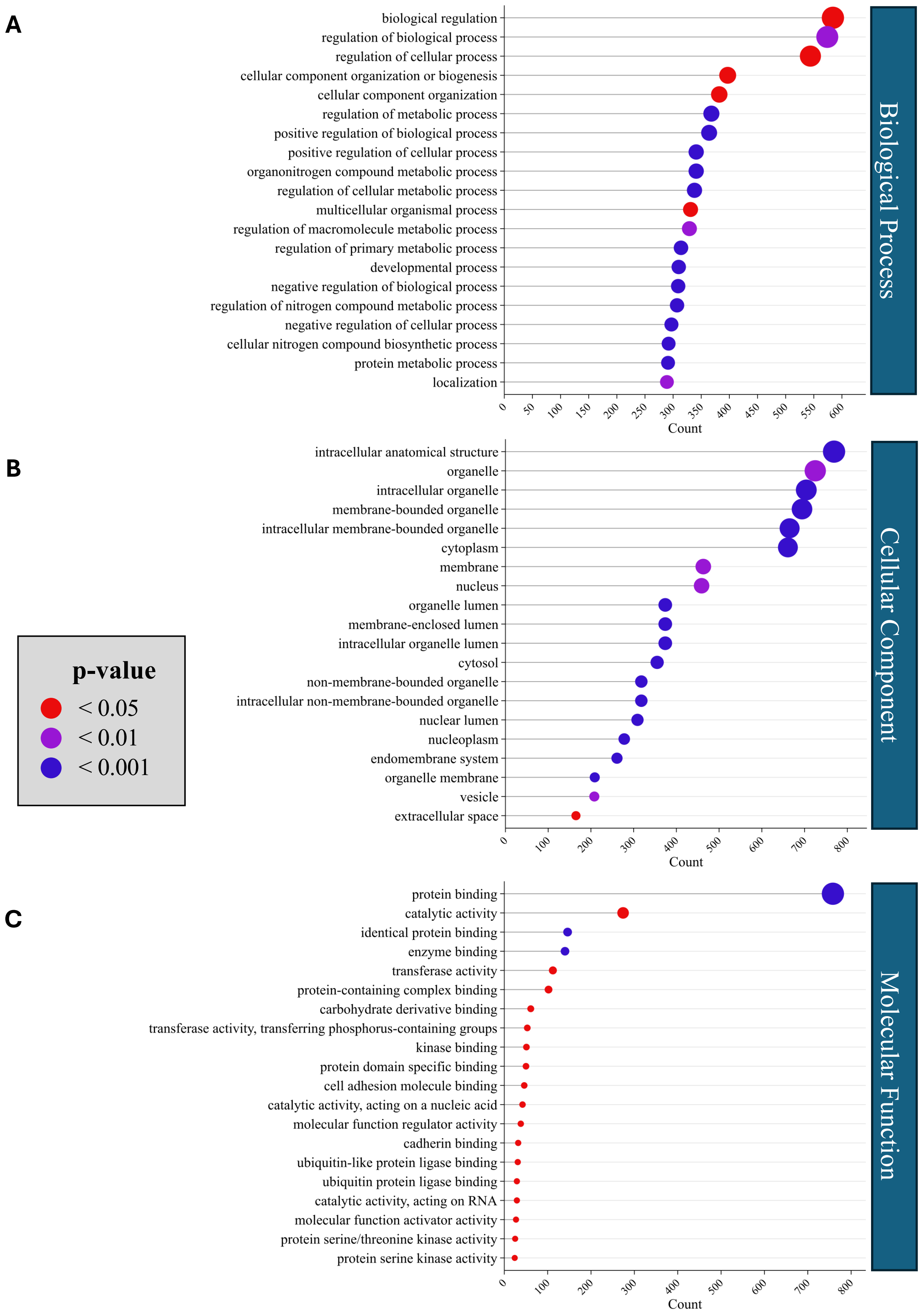


**Figure S11: Gene Ontology (GO) Enrichment Analysis of DEGs Uniquely Regulated by COX4I2 Knockdown**

The top 20 significantly enriched (adjusted p<0.05) GO terms of the 1440 genes uniquely differentially expressed between siControl and siCOX4I2 in the **A**) Biological Process, **B**) Cellular Component, and **C**) Molecular Function domains. The DEGs uniquely regulated by siCOX4I2 related to regulation of biological, cellular, and metabolic processes, as well as protein and enzyme binding GO pathways, with GO cellular components relating to organelles, membranes, nucleus, vesicle, and cytosol.

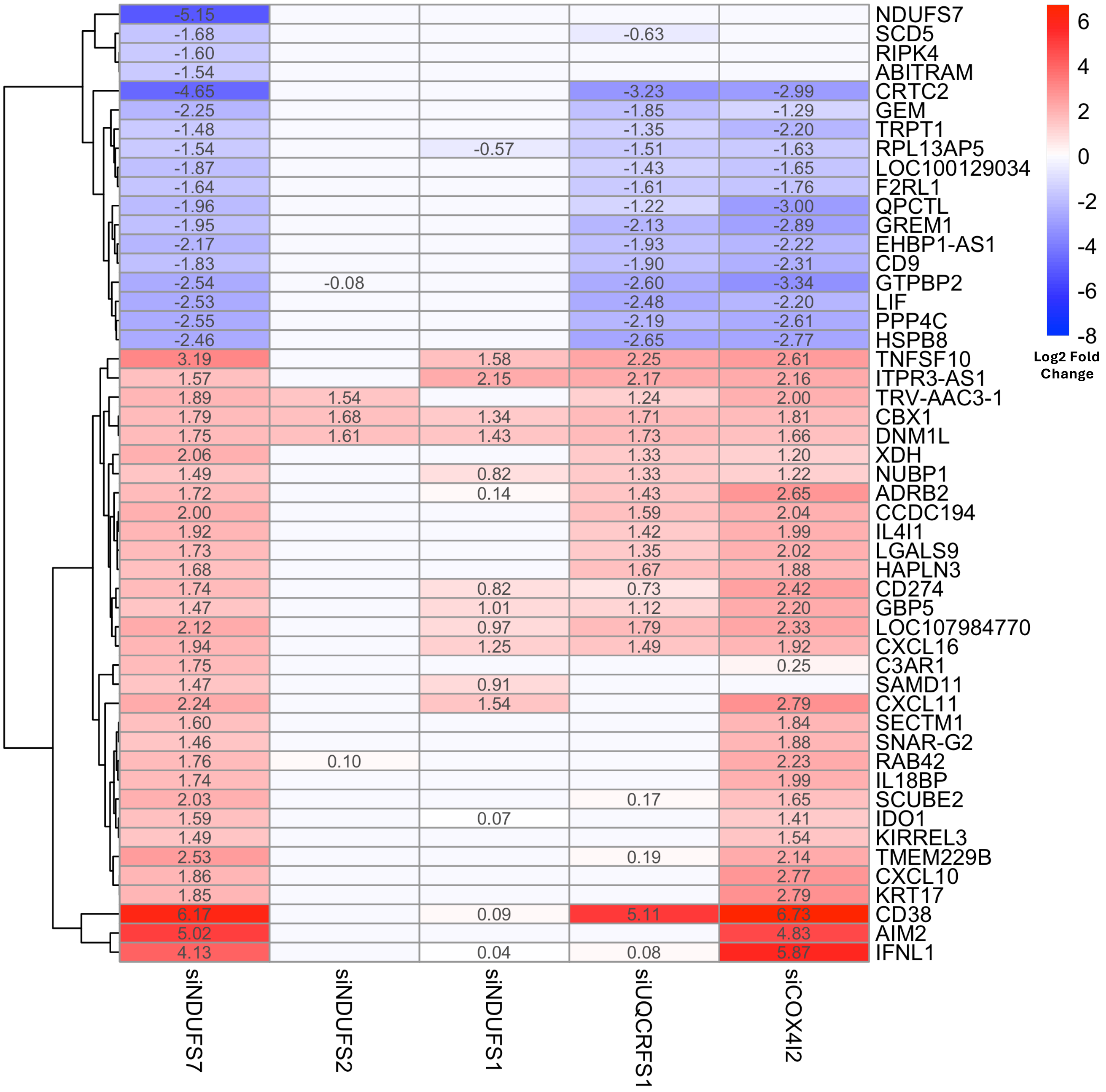


**Figure S12: DEGs with NDUFS7 Knockdown Reveals Distinct Gene Expression Pattern Among ETC Complex I Knockdowns**

Comparing the top 50 siNDUFS7 versus siControl DEGs to other selective knockdown conditions, there is little overlap in the effects of siNDUFS7 and the other ETC Complex I knockdowns, siNDUFS2 and siNDUFS1. There is minimal overlap in the genes most changed with NDUFS7 knockdown and the significant DEGs with knockdown of another subunit in the ubiquinone binding pocket, NDUFS2, demonstrating the differential effect of knocking down these neighbouring subunits. The genes most up- and downregulated by NDUFS7 knockdown are similarly affected by COX4I2 knockdown, with more than half of these also being similarly affected by UQCRFS1 knockdown.

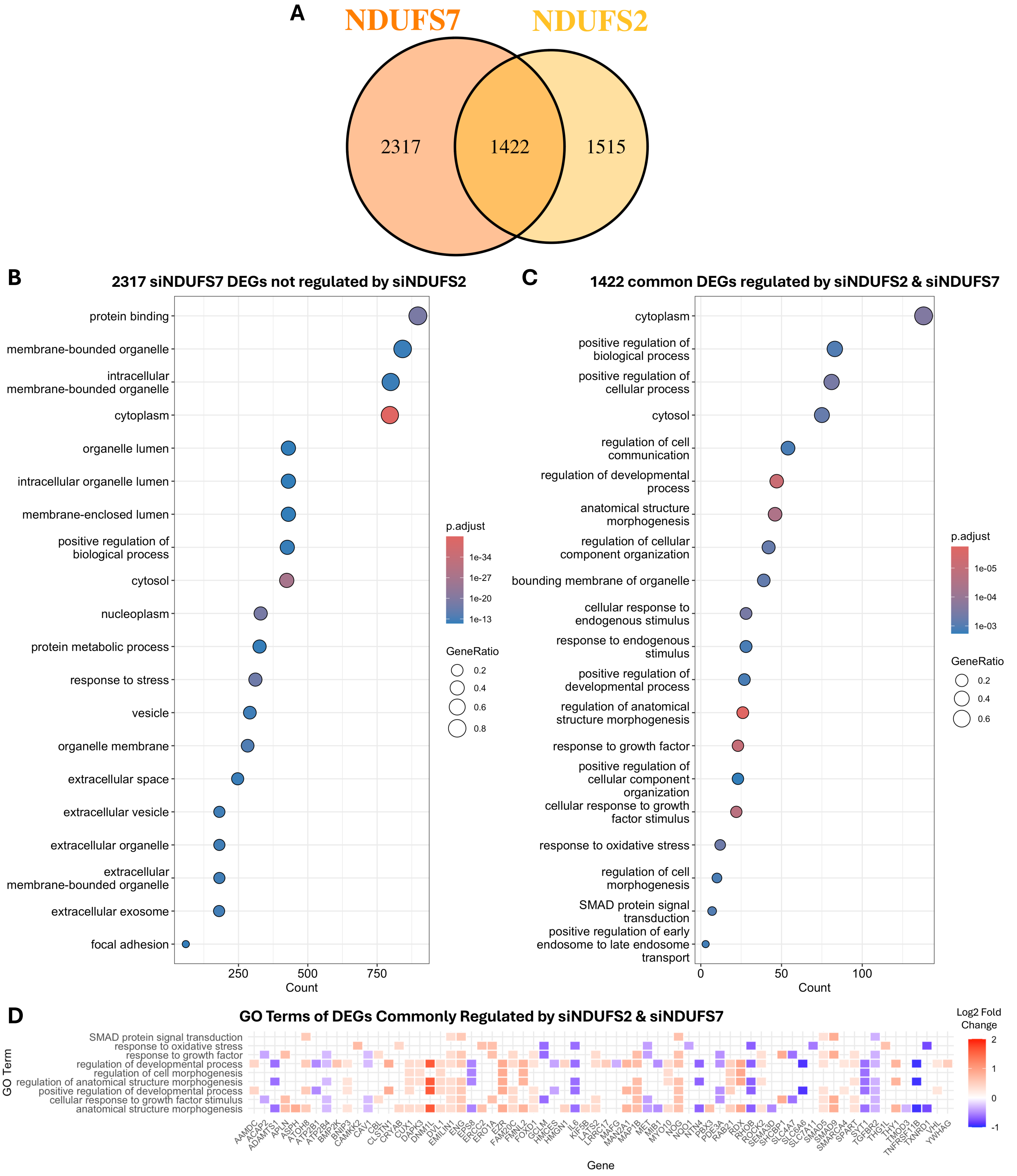


**Figure S13: Gene Ontology (GO) Enrichment Analysis of DEGs Uniquely and Commonly Regulated by Knockdown of Components of the Ubiquinone Binding Pocket**

**A**) Comparison of significant (adjusted p<0.05) DEGs between siControl and siNDUFS7 (left) and between siControl and siNDUFS2 (right). **B**) The top 20 significantly enriched (adjusted p<0.05) GO terms of the 2317 genes regulated by NDUFS7 knockdown but not NDUFS2 knockdown reveal enrichment of structural and vesicle/secretory pathways. **C**) The top 20 significantly enriched GO terms of the 1422 genes commonly regulated by NDUFS7 and NDUFS2 knockdowns include enrichment of pathways relating to response to cell stimuli, developmental processes, and response to oxidative stress. **D**) Heatmap of the DEGs within the growth, development, and oxidative stress pathways from the top 20 GO terms common between siNDUFS2 and siNDUFS7 in panel C (coloured by Log2 Fold Change of siControl vs siNDUFS2).

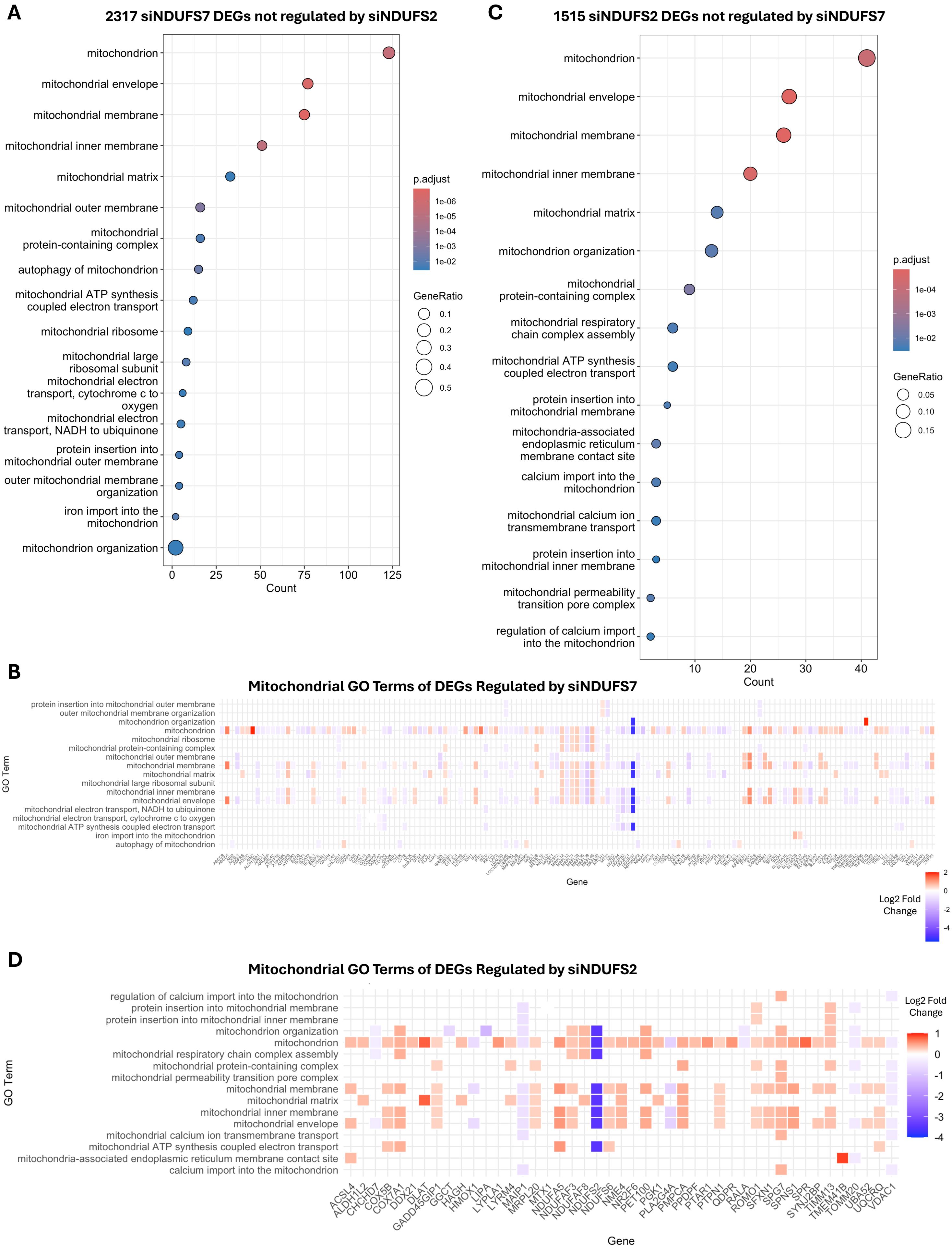
A
2317 siNDUFS7 DEGs not regulated by siNDUFS2
mitochondrion
mitochondrial envelope
mitochondrial membrane
mitochondrial inner membrane
mitochondrial matrix
mitochondrial outer membrane
mitochondrial protein-containing complex
autophagy of mitochondrion
mitochondrial ATP synthesis coupled electron transport
mitochondrial ribosome
mitochondrial large ribosomal subunit
mitochondrial electron transport, cytochrome c to oxygen
mitochondrial electron transport, NADH to ubiquinone
protein insertion into mitochondrial outer membrane
outer mitochondrial membrane organization
iron import into the mitochondrion
mitochondrion organization
0
25
50
75
100
125
Count
p.adjust
1e-06
1e-05
1e-04
1e-03
1e-02
GeneRatio
0.1
0.2
0.3
0.4
0.5
C
1515 siNDUFS2 DEGs not regulated by siNDUFS7
mitochondrion
mitochondrial envelope
mitochondrial membrane
mitochondrial inner membrane
mitochondrial matrix
mitochondrion organization
mitochondrial protein-containing complex
mitochondrial respiratory chain complex assembly
mitochondrial ATP synthesis coupled electron transport
protein insertion into mitochondrial membrane
mitochondria-associated endoplasmic reticulum membrane contact site
calcium import into the mitochondrion
mitochondrial calcium ion transmembrane transport
protein insertion into mitochondrial inner membrane
mitochondrial permeability transition pore complex
regulation of calcium import into the mitochondrion
10
20
30
40
Count
p.adjust
1e-04
1e-03
1e-02
GeneRatio
0.05
0.10
0.15
B
Mitochondrial GO Terms of DEGs Regulated by siNDUFS7
protein insertion into mitochondrial outer membrane
outer mitochondrial membrane organization
mitochondrion organization
mitochondrion
mitochondrial ribosome
mitochondrial protein-containing complex
mitochondrial outer membrane
mitochondrial membrane
mitochondrial matrix
mitochondrial large ribosomal subunit
mitochondrial inner membrane
mitochondrial envelope
mitochondrial electron transport, NADH to ubiquinone
mitochondrial electron transport, cytochrome c to oxygen
mitochondrial ATP synthesis coupled electron transport
iron import into the mitochondrion
autophagy of mitochondrion
GO Term
Gene
Log2 Fold Change
2
0
-2
-4
D
Mitochondrial GO Terms of DEGs Regulated by siNDUFS2
regulation of calcium import into the mitochondrion
protein insertion into mitochondrial membrane
protein insertion into mitochondrial inner membrane
mitochondrion organization
mitochondrion
mitochondrial respiratory chain complex assembly
mitochondrial protein-containing complex
mitochondrial permeability transition pore complex
mitochondrial membrane
mitochondrial matrix
mitochondrial inner membrane
mitochondrial envelope
mitochondrial calcium ion transmembrane transport
mitochondrial ATP synthesis coupled electron transport
mitochondria-associated endoplasmic reticulum membrane contact site
calcium import into the mitochondrion
GO Term
ACSL4
ALDH1L2
CHCHD7
COX5B
COX7A1
DDX21
DLAT
GADD45GIP1
GGCT
HAGH
HMOX1
LIPA
LYPLA1
LYRM4
MAIP1
MRPL20
MTX1
NDUFA5
NDUFAF3
NDUFAF8
NDUFS2
NDUFS6
NME4
NR2F6
PET100
PGK1
PLA2G4A
PMPCA
PPDPF
PTAR1
PTPN1
QDPR
RALA
ROMO1
SFXN1
SPG7
SPNS1
SPR
SYNJ2BP
TIMM13
TMEM41B
TOMM20
UBA52
UQCRQ
VDAC1
Gene
Log2 Fold Change
1
0
-1
-2
-3
-4

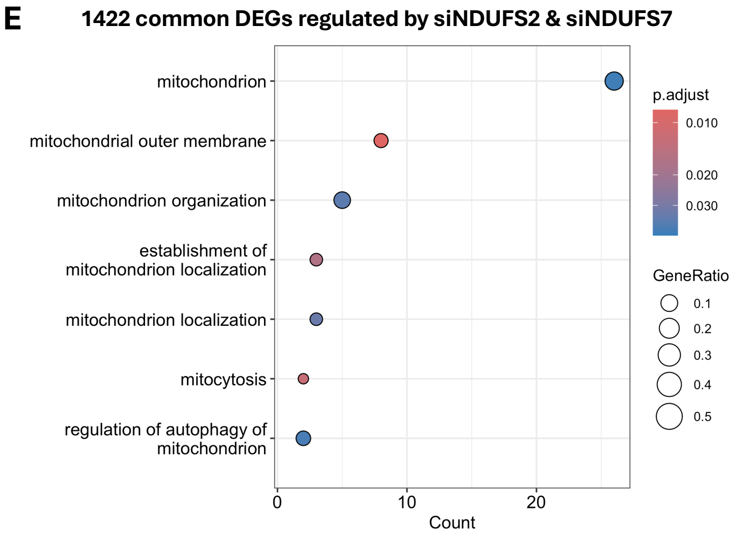


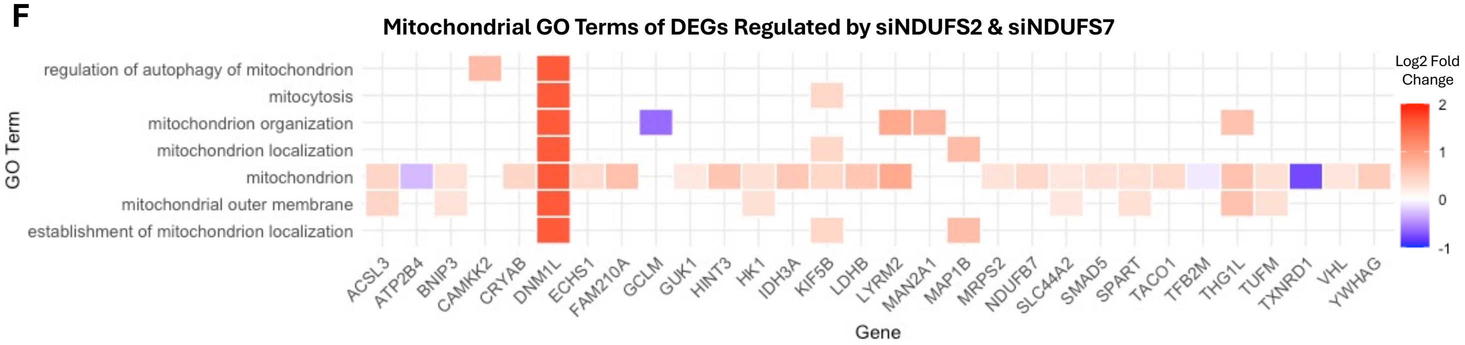


**Figure S14: Mitochondrial Gene Ontology (GO) Enrichment Analysis of DEGs Uniquely and Commonly Regulated by Knockdown of Components of the Ubiquinone Binding Pocket, NDUFS2 and NDUFS7**

**A**) Significantly enriched (adjusted $p<0.05$) mitochondrial GO terms for the 2317 genes regulated by NDUFS7 knockdown but not NDUFS2 knockdown, filtered with the search string "mitoc". **B**) The majority of the DEGs within the GO mitochondrial pathways (**S14A**) are downregulated by NDUFS7 knockdown. **C**) Significantly enriched mitochondrial GO terms for the 1515 genes regulated by NDUFS2 knockdown but not NDUFS7 knockdown, filtered with the search string "mitoc". **D**) Most of the DEGs within the GO mitochondrial pathways (**S14C**) are upregulated by NDUFS2 knockdown. **E**) Significantly enriched mitochondrial GO terms for the 1422 genes commonly regulated by NDUFS7 and NDUFS2, filtered with the search string "mitoc". There were fewer mitochondrial pathways enriched within the commonly regulated genes than the unique DEGs with either NDUFS2 or NDUFS7 knockdown. **F**) As with the genes uniquely regulated by siNDUFS2, most of the commonly regulated genes in the mitochondrial GO pathways (**S14E**) were upregulated by the knockdowns (heatmap coloured by Log2 Fold Change of siControl vs siNDUFS2).

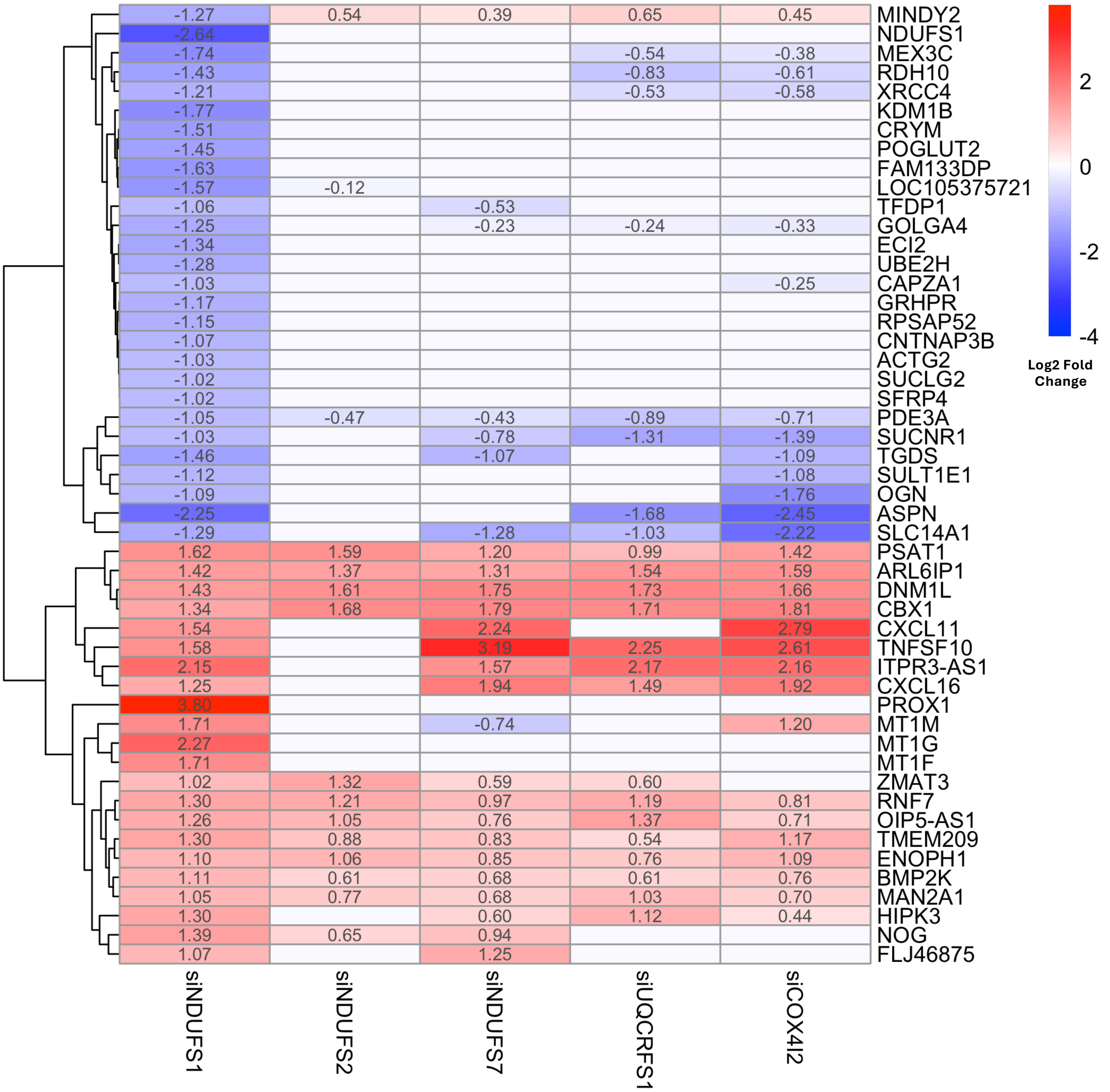


**Figure S15: DEGs with NDUFS1 Knockdown Reveals Distinct Gene Expression Pattern Among Selective ETC Knockdowns**

Comparing the top 50 siNDUFS1 versus siControl DEGs to other selective knockdown conditions, there is little overlap in the genes most downregulated by siNDUFS1 and the other ETC knockdowns. There is similar upregulation of some of the genes most upregulated by siNDUFS1 with the other knockdown conditions, with NDUFS2 knockdown having the least overlap of significant DEGs with genes most upregulated by siNDUFS1.

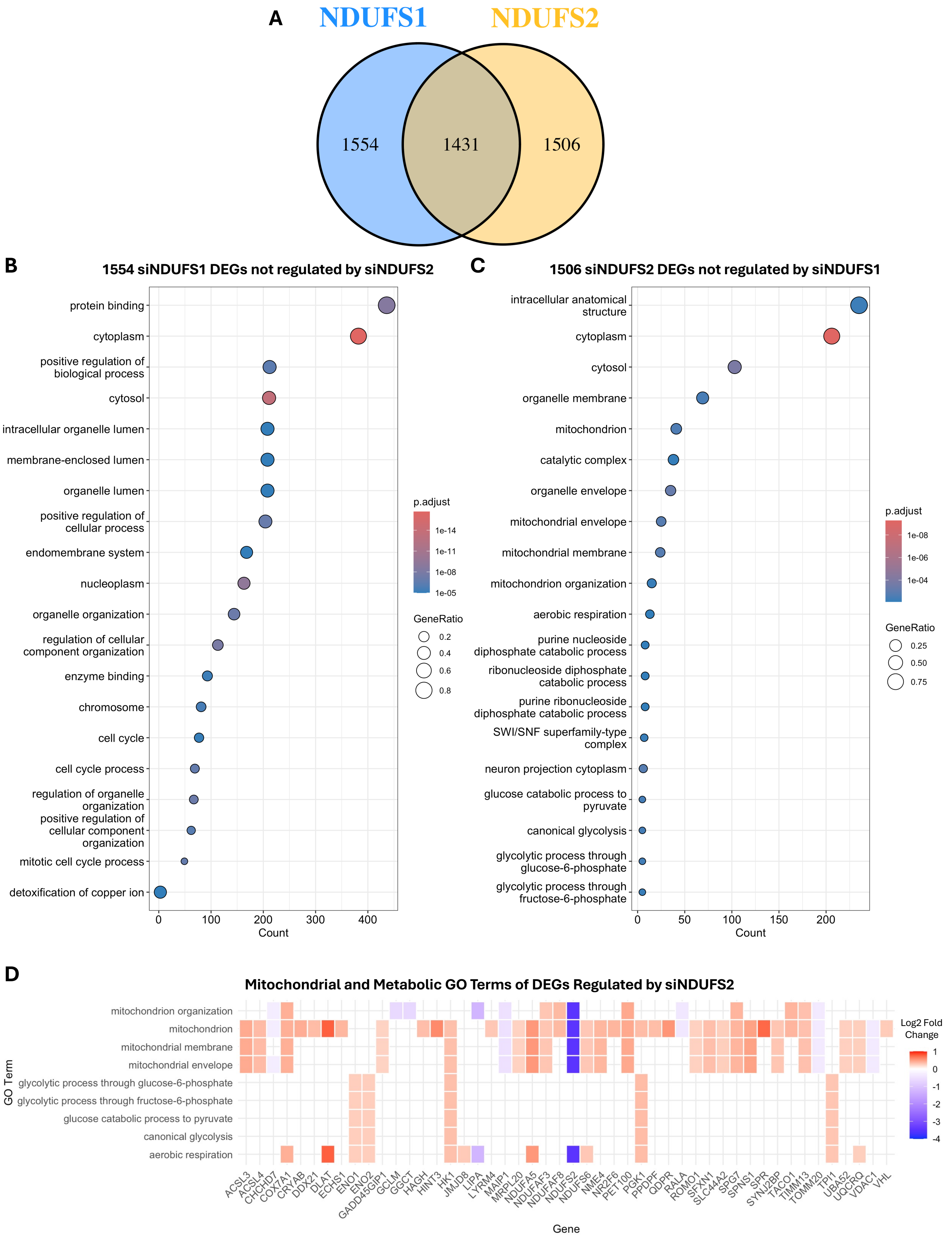

A
NDUFS1
NDUFS2
1554
1431
1506
B
1554 siNDUFS1 DEGs not regulated by siNDUFS2
protein binding
cytoplasm
positive regulation of biological process
cytosol
intracellular organelle lumen
membrane-enclosed lumen
organelle lumen
positive regulation of cellular process
endomembrane system
nucleoplasm
organelle organization
regulation of cellular component organization
enzyme binding
chromosome
cell cycle
cell cycle process
regulation of organelle organization
positive regulation of cellular component organization
mitotic cell cycle process
detoxification of copper ion
0
100
200
300
400
Count
p.adjust
1e-14
1e-11
1e-08
1e-05
GeneRatio
0.2
0.4
0.6
0.8
C
1506 siNDUFS2 DEGs not regulated by siNDUFS1
intracellular anatomical structure
cytoplasm
cytosol
organelle membrane
mitochondrion
catalytic complex
organelle envelope
mitochondrial envelope
mitochondrial membrane
mitochondrion organization
aerobic respiration
purine nucleoside diphosphate catabolic process
ribonucleoside diphosphate catabolic process
purine ribonucleoside diphosphate catabolic process
SWI/SNF superfamily-type complex
neuron projection cytoplasm
glucose catabolic process to pyruvate
canonical glycolysis
glycolytic process through glucose-6-phosphate
glycolytic process through fructose-6-phosphate
0
50
100
150
200
Count
p.adjust
1e-08
1e-06
1e-04
GeneRatio
0.25
0.50
0.75
D
Mitochondrial and Metabolic GO Terms of DEGs Regulated by siNDUFS2
GO Term
mitochondrion organization
mitochondrion
mitochondrial membrane
mitochondrial envelope
glycolytic process through glucose-6-phosphate
glycolytic process through fructose-6-phosphate
glucose catabolic process to pyruvate
canonical glycolysis
aerobic respiration
ACSL3
ACSL4
CHCHD7
COX7A1
CRYAB
DDX21
DLAT
ECHS1
ENO1
ENO2
GADD45GIP1
GCLM
GGCT
HAGH
HINT3
HK1
JMJD8
LIPA
LYRM4
MAIP1
MRPL20
NDUFA5
NDUFAF3
NDUFAF8
NDUFS2
NDUFS6
NME4
NR2F6
PET100
PGK1
PPDPF
QDPR
RALA
ROMO1
SFXN1
SLC44A2
SPG7
SPNS1
SPR
SYNJ2BP
TACO1
TIMM13
TOMM20
TPI1
UBA52
UQCRQ
VDAC1
VHL
Gene
Log2 Fold Change
1
0
-1
-2
-3
-4

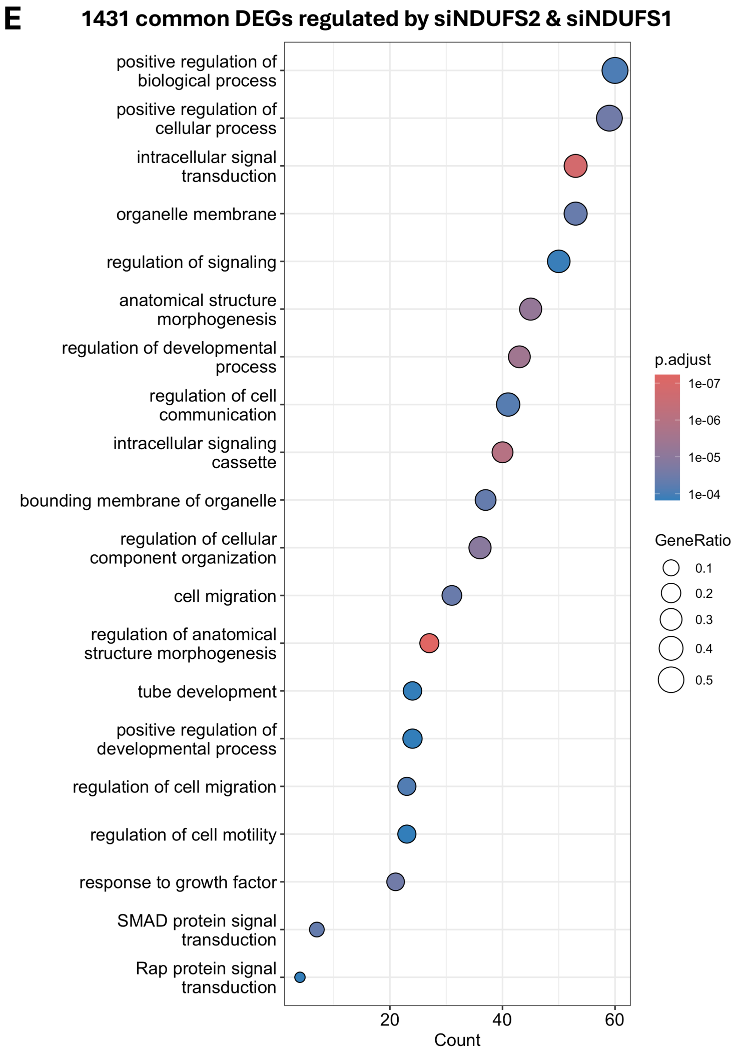


**Figure S16: Gene Ontology (GO) Enrichment Analysis of DEGs Uniquely and Commonly Regulated by Knockdown of Complex I Subunits NDUFS2 and NDUFS1**

**A**) Comparison of significant (adjusted p<0.05) DEGs between siControl and siNDUFS1 (left) and between siControl and siNDUFS2 (right), providing a comparison of the effects of knocking down NDUFS2 in the ubiquinone binding pocket and NDUFS1 in the peripheral arm of Complex I. **B**) The top 20 significantly enriched (adjusted p<0.05) GO terms of the 1554 genes regulated by NDUFS1 knockdown but not NDUFS2 knockdown reveal enrichment of organellar organization and cell cycle pathways. **C**) The top 20 significantly enriched GO terms of the 1506 genes regulated by NDUFS2 knockdown but not NDUFS1 knockdown reveal enrichment of mitochondrial and metabolic pathways. **D**) A heatmap of the DEGs within the mitochondrial pathways in the top 20 GO terms enriched with NDUFS2 knockdown (from **S16C**) shows the majority of mitochondrial and respiratory genes are upregulated by siNDUFS2 relative to siControl (coloured by Log2 Fold Change of siControl vs siNDUFS2). **E**) The top 20 significantly enriched GO terms of the 1431 genes commonly regulated by NDUFS1 and NDUFS2 knockdowns include enrichment of pathways relating to cell signalling and migration.

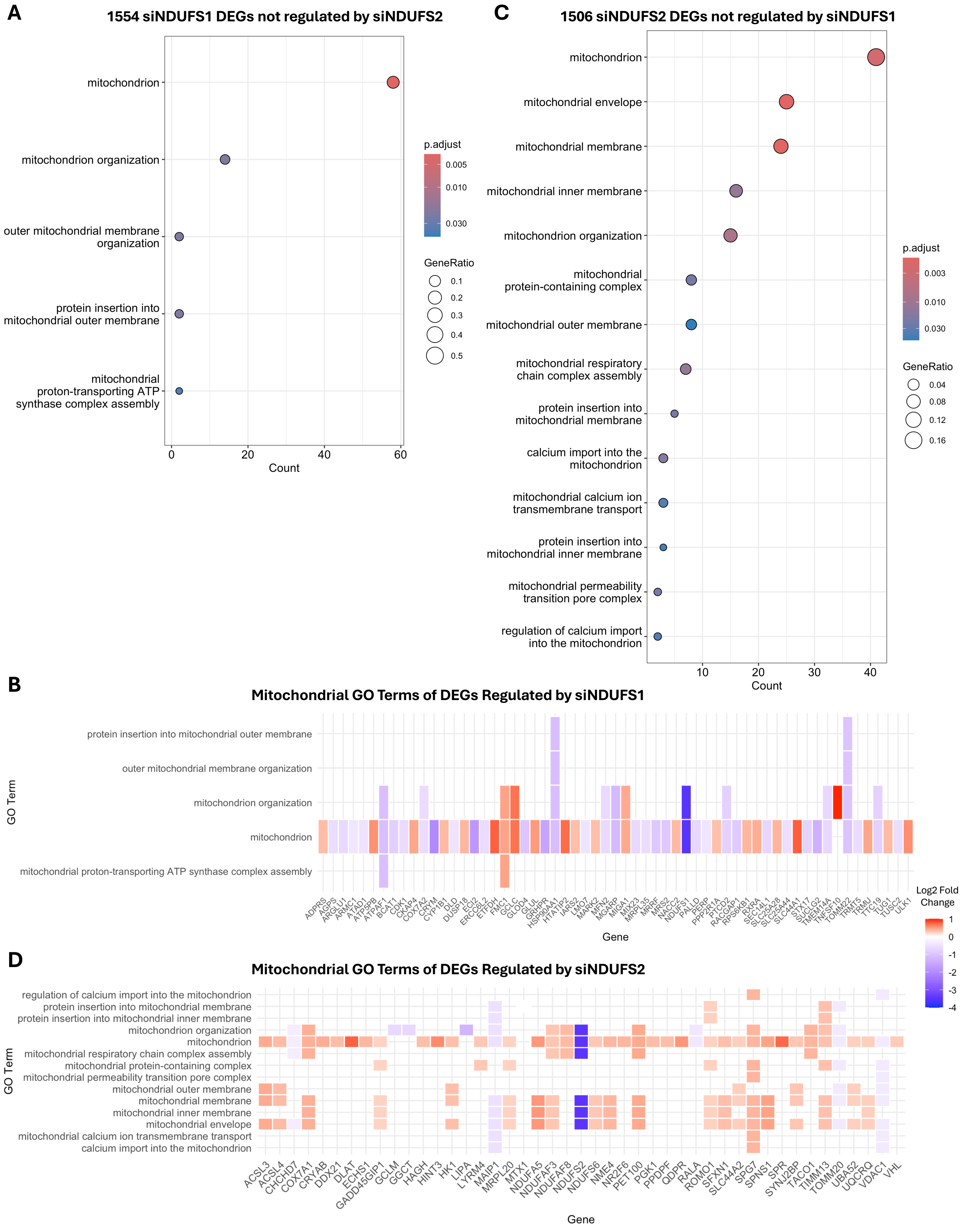
A
1554 siNDUFS1 DEGs not regulated by siNDUFS2
mitochondrion
mitochondrion organization
outer mitochondrial membrane organization
protein insertion into mitochondrial outer membrane
mitochondrial proton-transporting ATP synthase complex assembly
0
20
40
60
Count
p.adjust
0.005
0.010
0.030
GeneRatio
0.1
0.2
0.3
0.4
0.5
C
1506 siNDUFS2 DEGs not regulated by siNDUFS1
mitochondrion
mitochondrial envelope
mitochondrial membrane
mitochondrial inner membrane
mitochondrion organization
mitochondrial protein-containing complex
mitochondrial outer membrane
mitochondrial respiratory chain complex assembly
protein insertion into mitochondrial membrane
calcium import into the mitochondrion
mitochondrial calcium ion transmembrane transport
protein insertion into mitochondrial inner membrane
mitochondrial permeability transition pore complex
regulation of calcium import into the mitochondrion
10
20
30
40
Count
p.adjust
0.003
0.010
0.030
GeneRatio
0.04
0.08
0.12
0.16
B
Mitochondrial GO Terms of DEGs Regulated by siNDUFS1
GO Term
protein insertion into mitochondrial outer membrane
outer mitochondrial membrane organization
mitochondrion organization
mitochondrion
mitochondrial proton-transporting ATP synthase complex assembly
ADPRS AGPS ARGLU1 ARMC1 ATAD1 ATP5PB ATPAF1 BCAT1 CDK1 CKAP4 COX7A2 CRYM CYP1B1 DLD DUSP18 ECI2 ERCC6L2 ETFDH FMC1 GCLC GLOD4 GLUL GRHPR HSP90AA1 HTATIP2 IARS2 LMO7 MARK2 MFN2 MGARP MIGA1 MIX23 MRPL35 MRRF MRS2 NACC2 NDUFS1 PALLD PERP PPP2R1A PTCD2 RACGAP1 RPS6KB1 RXRA SEC14L1 SLC25A28 SLC25A44 SLC44A1 STX17 SUCLG2 TMEM14A TNFSF10 TOMM22 TRMT5 TRMU TTC19 TUG1 TUSC2 ULK1
Gene
Log2 Fold Change
1
0
-1
-2
-3
-4
D
Mitochondrial GO Terms of DEGs Regulated by siNDUFS2
GO Term
regulation of calcium import into the mitochondrion
protein insertion into mitochondrial membrane
protein insertion into mitochondrial inner membrane
mitochondrion organization
mitochondrion
mitochondrial respiratory chain complex assembly
mitochondrial protein-containing complex
mitochondrial permeability transition pore complex
mitochondrial outer membrane
mitochondrial membrane
mitochondrial inner membrane
mitochondrial envelope
mitochondrial calcium ion transmembrane transport
calcium import into the mitochondrion
ACSL3 ACSL4 CHCHD7 COX7A1 CRYAB DDX21 DLAT ECHS1 GADD45GIP1 GCLM GGCT HAGH HINT3 HK1 LIPA LYRM4 MAIP1 MRPL20 MTX1 NDUFA5 NDUFAF3 NDUFAF8 NDUFS2 NDUFS6 NME4 NR2F6 PET100 PGK1 PPDPF QDPR RALA ROMO1 SFXN1 SLC44A2 SPG7 SPNS1 SPR SYNJ2BP TACO1 TIMM13 TOMM20 UBA52 UQCRQ VDAC1 VHL
Gene

E

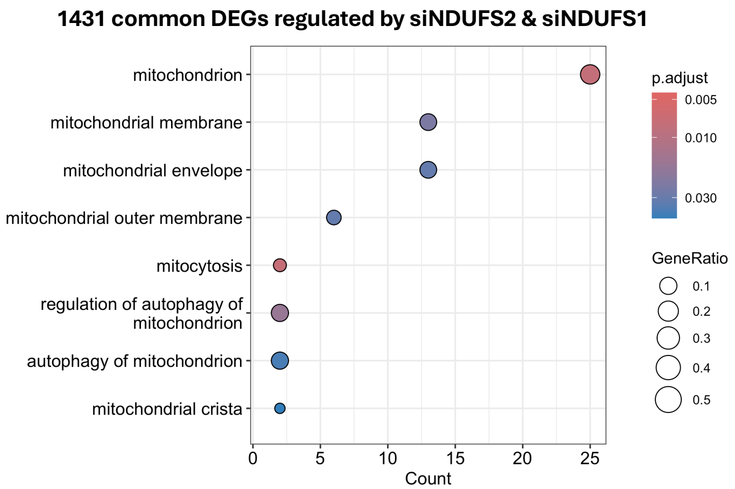


F

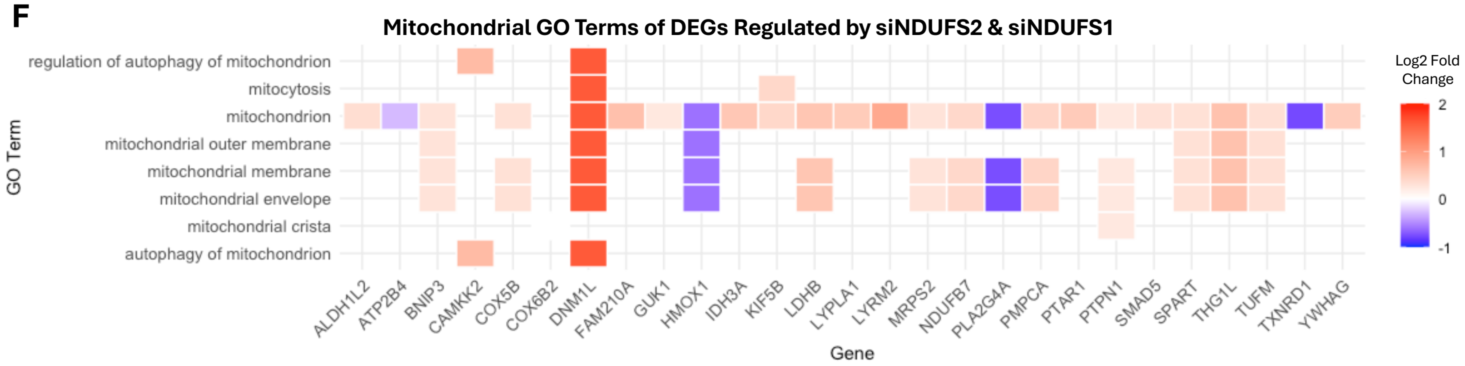


**Figure S17: Mitochondrial Gene Ontology (GO) Enrichment of DEGs Uniquely and Commonly Regulated by Knockdown of Complex I Subunits NDUFS2 and NDUFS1**

Comparison of the mitochondrial gene expression changes between knocking down NDUFS2 in the ubiquinone binding pocket and NDUFS1 in the peripheral arm of Complex I. **A**) Significantly enriched (adjusted $p<0.05$) mitochondrial GO terms for the 1554 genes regulated by NDUFS1 knockdown but not NDUFS2 knockdown, filtered with the search string "mitoc". There were few enriched mitochondrial GO terms with NDUFS1 knockdown and **B**) most of the genes in the mitochondrial GO pathways are downregulated by siNDUFS1. **C**) Significantly enriched mitochondrial GO terms for the 1506 genes regulated by NDUFS2 knockdown but not NDUFS1 knockdown, filtered with the search string "mitoc". There were many more mitochondrial GO terms enriched with NDUFS2 knockdown, with **D**) most of the genes in the mitochondrial GO pathways being upregulated by siNDUFS2. **E**) Significantly enriched mitochondrial GO terms for the 1431 genes commonly regulated by NDUFS1 and NDUFS2 knockdowns, filtered with the search string "mitoc". There were more enriched mitochondrial GO terms than for the genes uniquely regulated by siNDUFS1, though there were fewer mitochondrial genes captured in these GO pathways. **F**) As with the genes uniquely regulated by siNDUFS2, most of the genes commonly regulated by siNDUFS1 and siNDUFS2 in the mitochondrial GO pathways were upregulated by the knockdowns (heatmap coloured by Log2 Fold Change of siControl vs siNDUFS2).
